# Synthetic lethality across normal tissues is strongly associated with cancer risk, onset, and tumor suppressor specificity

**DOI:** 10.1101/818922

**Authors:** Kuoyuan Cheng, Nishanth Ulhas Nair, Joo Sang Lee, Eytan Ruppin

## Abstract

Various characteristics of cancers exhibit tissue-specificity, including lifetime cancer risk, onset age and cancer driver genes. Previously, the large variation in cancer risk across human tissues was found to strongly correlate with the number of stem cell divisions and abnormal DNA methylation levels occurring in them. Here we study the role of another potentially important factor, synthetic lethality, in cancer risk. Analyzing transcriptomics data in the GTEx compendium we quantify the extent of co-inactivation of cancer synthetic lethal (cSL) gene pairs in *normal* tissues and find that normal tissues with more down-regulated cSL gene pairs have lower and delayed cancer risk. We also show that the tissue-specificity of numerous tumor suppressor genes is strongly associated with the expression of their cSL partner genes in the corresponding normal tissues. Overall, our findings uncover the role of synthetic lethality as a novel important factor involved in tumorigenesis.

## Introduction

Cancers of different human tissues have markedly different molecular, phenotypic and epidemiological characteristics, known as the tissue-specificity in cancer. Various aspects of this intriguing phenomenon include a considerable variation in lifetime cancer risk, cancer onset age and the genes driving the cancer across tissue types. The variation in lifetime cancer risk is known to span several orders of magnitude (1,2). Such variation cannot be fully explained by the difference in exposure to carcinogens or hereditary factors, and has been shown to strongly correlate with differences in the number of lifetime stem cell divisions (NSCD) estimated across tissues (2,3). As claimed by Tomasetti and Vogelstein, 2015 (2), these findings are consistent with the notion that tissue stem cell divisions can propagate mutations caused either by environmental carcinogens or random replication error (4). Additionally, the importance of epigenetic factors in carcinogenesis has long been recognized (5), and Klutstein *et al*. have recently reported that the levels of abnormal CpG island DNA methylation (LADM) across tissues is highly correlated with their cancer risk (6). Although both global (e.g. smoking and obesity) and various cancer type-specific (e.g. HCV infection for liver cancer) risk factors are well-known (7), no factors other than NSCD and LADM have been reported to date to explain the *across-tissue variance* in lifetime cancer risk.

Besides lifetime cancer risk, cancer onset age, as measured by the median age at diagnosis, also varies among adult cancers (1). Although most cancers typically manifest later in life (over 40 years old (1,8)), some such as testicular cancer often have earlier onset (1). Many tumor suppressor genes and oncogenes are also tissue specific (9–11). For example, mutations in the tumor suppressor gene *BRCA1* are predominantly known to drive the development of breast and ovarian cancer, but rarely other cancer types (12). In general, factors explaining the overall tissue-specificity in cancer could be tissue-intrinsic (10,13), and their elucidation can further advance our understanding of the forces driving carcinogenesis.

*Synthetic lethality/sickness* (SL) is a well-known type of genetic interaction, conceptualized as cell death or reduced cell viability that occurs under the combined inactivation of two genes, but not under the inactivation of either gene alone. The phenomenon of SL interactions was first recorded in *Drosophila* (14) and then in *Saccharomyces cerevisiae* (15). In recent years, much effort has been made to identify SL interactions specifically in cancer, since targeting these cancer SLs (cSLs) has been recognized as a highly valuable approach for cancer treatment (16–19). The effect of cSL on cancer cell viability has led us to investigate whether it plays an additional role even before tumors manifest, i.e. during carcinogenesis. In this study we quantify the level of cSL gene pair co-inactivation in normal (non-cancerous) human tissue as a measure of resistance to cancer development (termed cSL load, explained in detail below). We show that cSL load can explain a considerable level of the variation in cancer risk and cancer onset age across human tissues, as well as the tissue-specificity of some tumor suppressor genes. Taken together, our findings support the importance of synthetic lethality in impeding tumorigenesis across human tissues.

## Results

### Computing cSL load in normal human tissues

To study the potential effects of cancer synthetic lethality (cSL) in *normal, non-cancerous* tissues, we define a measure called cSL load, which quantifies the level of cSL gene pair co-inactivation based on gene expression of *normal* human tissues from the GTEx dataset (20). Specifically, we used a recently published reference set of genome-wide cSLs that are common to many cancer types, identified from both in vitro and TCGA cancer patient data (21) via the ISLE method (22,23) (Table S1a). For each GTEx normal tissue sample, we computed the cSL load as the fraction of cSL gene pairs (among all the genome-wide cSLs) that have both genes lowly expressed in that sample (Methods; illustrated in **Fig. 1**). We further defined tissue cSL load (TCL) as the median cSL load value across all samples of each tissue type in GTEx (Methods, Table S2a). We then proceed to test our hypothesis that TCL can be a measure of the level of resistance to cancer development intrinsic to each human tissue (outlined in **Fig. 1**).

**Figure 1.**
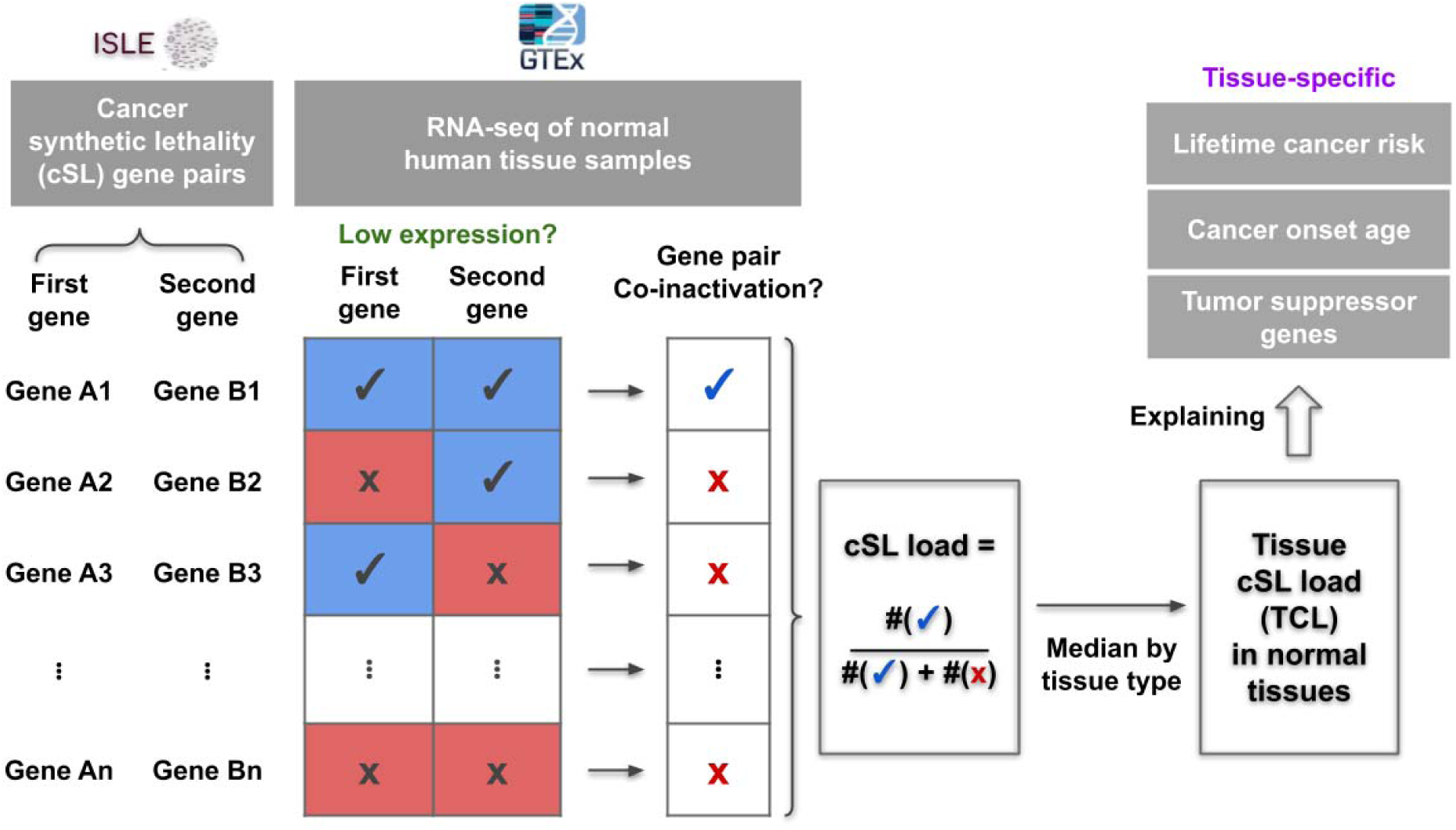
An illustration of the computation of cancer synthetic lethality (cSL) load for each sample and each tissue type (i.e. tissue cSL load, TCL), as well as an outline of this study where we attempted to explain the tissue-specific lifetime cancer risk, cancer onset age, and tumor suppressor genes using TCL. See main text and Methods for details.

### Tissue cSL load in normal tissues is inversely correlated with their lifetime cancer risk

Synthetic lethality is widely known to be context-specific across species, tissue types, and cellular conditions (24). In theory, a cancer-specific cSL gene pair can be co-inactivated in the normal tissue without reducing normal cell fitness, while conferring resistance to the emergence of malignantly transformed cells due to the lethal effect specifically on the cancer cells. Different normal tissues can have varied TCLs (representing the levels of cSL gene pair co-inactivations) as a result of their specific gene expression profiles, and we hypothesized that normal tissues with higher TCLs should have lower cancer risk, as transforming cancerous cells in these tissues will face higher cSL-mediated vulnerability and lethality. To test this hypothesis, we obtained data on the tissue-specific lifetime cancer risk in humans (Methods) and correlated that with the TCL values computed for the different tissue types. We find a strong negative correlation between the TCL (computed from older-aged GTEx samples, age ≥ 50 years) and lifetime cancer risk across normal tissues (Spearman’s ρ = −0.664, P = 1.59e-4, **Fig. 2a**, Table S2a). This correlation is robust, as comparable results are obtained when this analysis is carried out in various ways (e.g. different cutoffs for low expression of genes, different cSL network sizes, different cancer type-normal tissue mappings, etc., Fig. S1, Supp. Note). Notably, the cSL load varies with age due to age-related gene expression changes, and the correlation with lifetime cancer risk is not found when the TCL is computed on samples from the young population (20 ≤ age < 50 years, Spearman’s ρ = −0.0251, P = 0.901, Fig. S2a); this is consistent with the observation that lifetime cancer risk is mostly contributed by cancers occurring in older populations (1). Importantly, we still see a marked negative correlation between TCL and lifetime cancer risk when analyzing samples from all age groups together (Spearman’s ρ = −0.49, P = 0.01, Fig. S2b). Repeating these analyses using different control gene-pairs, including (i) random gene pairs; (ii) shuffled cSL gene pairs; and (iii) degree-preserving randomized cSL network (same size as the actual cSL network, Supp. Note) results in significantly weaker correlations (empirical P < 0.001, Fig. S3a-c, Supp. Note), confirming that the associations found with cancer risk results from a cSL-specific effect.

**Figure 2.**
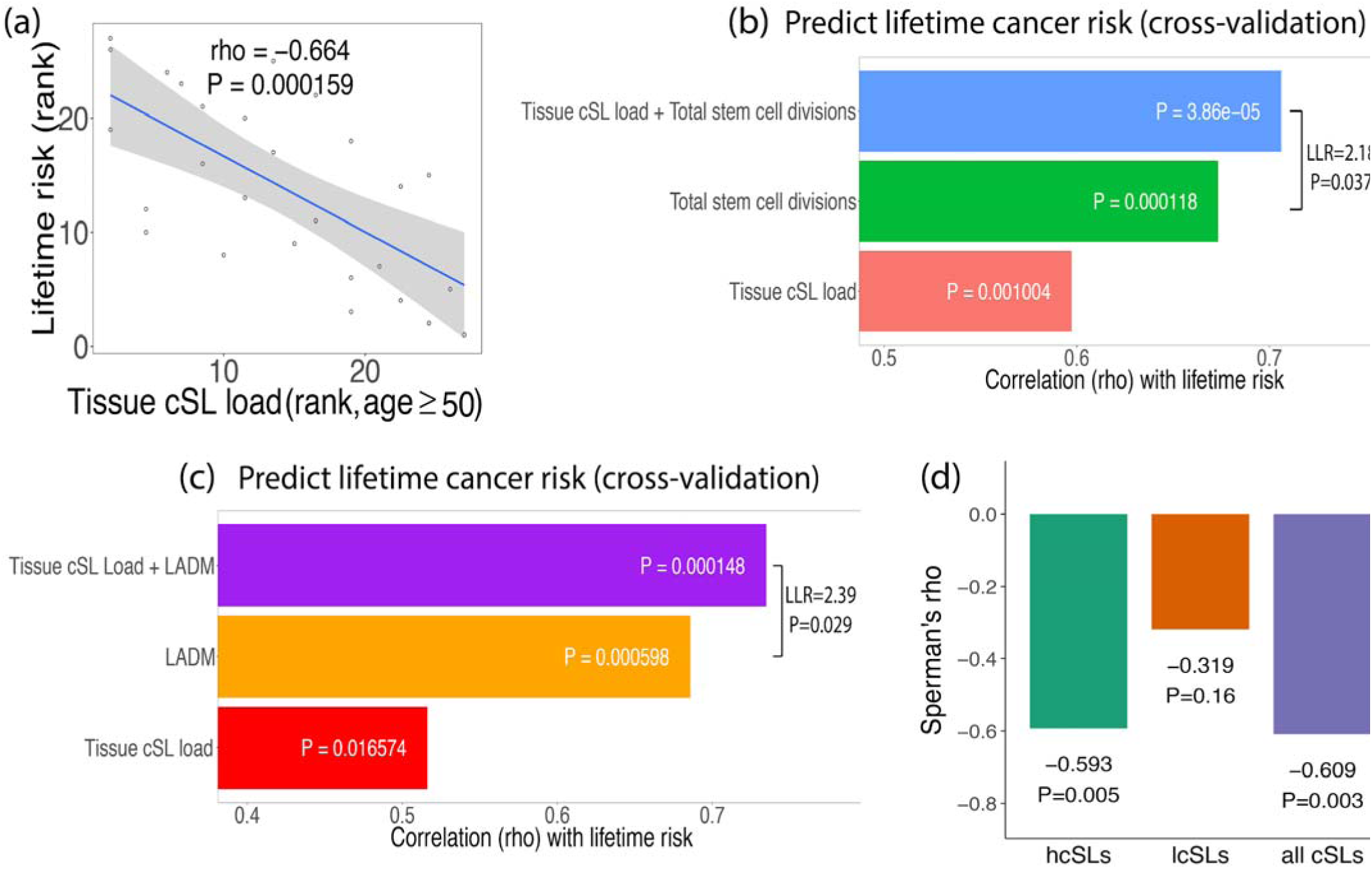
**(a)** Scatter plot showing the Spearman’s correlations between lifetime cancer risk and tissue cSL load (TCL) computed for the older population (age ≥ 50 years). (Ranked values are used as lifetime cancer risk spans several orders of magnitude.) **(b)** Lifetime cancer risks across tissues were predicted using linear models (under cross-validation) containing different sets of explanatory variables: (1) TCL only; (2) the number of stem cell divisions (NCSD) only; and (3) TCL and NSCD (27 data points). The prediction accuracy is measured by the Spearman’s ρ (rho), shown by the bar plots. The result of a likelihood ratio test between model (2) and model (3) is also displayed. **(c)** A similar bar plot as in (b) comparing the predictive models for cancer risk involving the variables: (1) TCL only; (2) the level of abnormal DNA methylation (LADM) only; and (2) TCL and LADM combined (21 data points only due to the smaller set of LADM data). A model containing all the three variables does not increase the prediction power (Spearman’s ρ = 0.77 under cross-validation) and is not shown. **(d)** Bar plot showing the correlations between lifetime cancer risk with TCLs computed (age ≥ 50 years) using subsets of cSLs — highly specific cSLs (hcSLs), lowly specific cSLs (lcSLs), and all cSLs. Spearman’s ρ and p-values are shown. The hcSLs and lcSLs are identified using data of matched TCGA cancer types and GTEx normal tissues (Methods), which corresponds to only a subset of tissue types. To facilitate comparison, here the correlation for all cSLs was also computed for the same subset of tissues, and therefore the resulting correlation coefficient is different from that in (a).

While the randomized cSL networks used in the control tests described above provide significantly weaker correlations with cancer risk than those observed with cSLs, many of these correlations are still significant by themselves (Fig. S3b,c). This suggests that there may be a possible association between the expression of single genes in the cSL network (*cSL genes*) and cancer risk. To investigate this, we computed the *tissue cSL single-gene load* (*SGL*, the fraction of lowly expressed cSL genes) for each tissue (Methods). Indeed, we do find a significant negative correlation between tissue SGL levels and cancer risk (Spearman’s ρ = −0.49, P = 0.01, Fig. S3d, Supp. Note). This correlation vanishes when we use random sets of single genes (Fig. S3f). However, after controlling for the single-gene effect, the partial correlation between tissue cSL load and cancer risk is still highly significant (Spearman’s rho = −0.69, P = 6.10e-5, Fig. S3g), pointing to the dominant role of the SL genetic interaction effect (Supp. Note).

### Tissue cSL load predicts lifetime cancer risk across tissues in addition to the number of tissue stem cell divisions and abnormal DNA methylation levels

We next compared the predictive power of TCL to those obtained with the previously reported measures of NSCD (2,3) and LADM (6), using the set of GTEx tissue types investigated here (Methods). We first confirmed the strong correlations of NSCD and LADM with tissue lifetime cancer risk in our specific dataset (Spearman’s ρ = 0.72 and 0.74, P = 2.6e-5 and 1.3e-4, respectively, Fig. S4). These correlations are stronger than the one we reported above between TCL and cancer risk. However, adding TCL to either NSCD or LADM in linear regression models leads to enhanced predictive models of cancer risk compared to those obtained with NSCD or LADM alone (log-likelihood ratio = 2.18 and 2.39, P = 0.037 and 0.029, respectively). Furthermore, adding TCL to each of these factors increases their prediction accuracy under cross-validation (Spearman’s ρ’s from 0.67 and 0.69 with NSCD and LADM alone to 0.71 and 0.77, respectively, **Fig. 2b,c**). LADM and NSCD are significantly correlated (Spearman’s ρ = 0.66, P = 0.02), while the TCL correlates only in a borderline significant manner with either NSCD (Spearman’s ρ = −0.57, P = 0.06) or LADM (Spearman’s ρ = −0.52, P = 0.08). Taken together, these observations support the hypothesis that TCL is associated with tissue cancer risk, with a partially independent role from either NSCD or LADM.

### cSL pairs that are more specific to cancer are more predictive of cancer risk in normal tissues

We have shown results that support the role of TCL in impeding cancer development, and we reason that such an effect is dependent on the notion that many of the cSLs are specific to cancer while having weaker or no lethal effects in normal tissues. Indeed, we tested and found that the co-inactivation of cSL gene pairs is under much weaker negative selection in GTEx normal tissues vs matched TCGA cancers (Wilcoxon rank-sum test P = 2.93e-6, Fig. S5a, also shown using cross-validation, Supp. Note). Moreover, we hypothesize that those cSLs with the highest specificity to cancer (i.e. with strongest SL effect in cancer and no or weakest effect on normal cells) should have the strongest effect on cancer development. To test this, we identified the subset of such cSLs (termed “highly specific cSLs” or “hcSLs”) as well as those with the lowest specificity to cancer (termed “lowly specific cSLs” or “lcSLs”; Methods), and re-computed the TCLs of all normal GTEx tissues using these two cSL subsets, respectively. Indeed, the TCLs computed from the hcSLs correlate much stronger with cancer lifetime risk than those computed from the lcSLs (Spearman’s ρ = −0.593 vs - 0.319, Fig. 2d), testifying that these cSLs with high functional specificity to cancer are truly relevant to carcinogenesis. These hcSLs are enriched for cell cycle, DNA damage response and immune-related genes (FDR < 0.05, Table S5, Methods), which are known to play key roles in tumorigenesis.

### Higher tissue cSL load in the younger population is associated with delayed cancer onset

We have thus established that TCL in the older population is inversely correlated with lifetime cancer risk across tissues. We next hypothesized that higher cSL load in a given normal tissue in the young population may delay cancer onset, which typically occurs later (age > 40 years (1)). To test this, we use the median age at cancer diagnosis (1) of a certain tissue as its cancer onset age (Table S3, Methods). We find that the TCL values (for age ≤ 40 years) are indeed markedly correlated with cancer onset age (Spearman’s ρ = 0.502, P = 0.011, **Fig. 3a**). This result is again robust to variations in our methods to compute TCL and cancer onset age (Fig. S6, Table S3, Supp. Note). We note that the cancer onset age is not significantly correlated with lifetime cancer risk (Spearman’s ρ = 0.279, P = 0.28).

**Figure 3.**
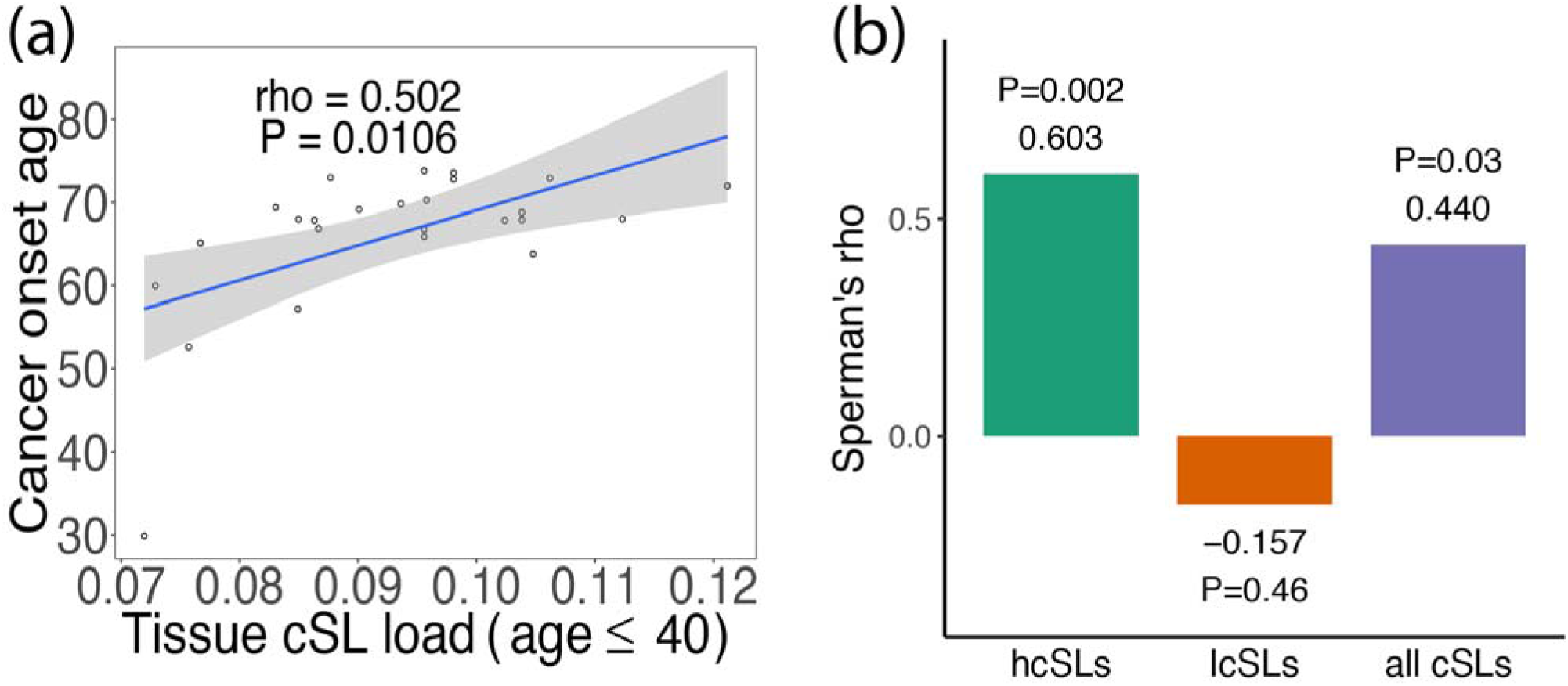
**(a)** Scatter plot showing the Spearman’s correlations between cancer onset age and TCL (age ≤ 40 years). **(b)** Bar plot showing the correlations between cancer onset age with TCLs computed (age ≤ 40 years) using subsets of cSLs — highly specific cSLs (hcSL), lowly specific cSL (lcSL), and all cSLs. Spearman’s ρ and p-values are shown. As in Fig. 2d, this analysis was done for a subset of GTEx normal tissues for which we had matched TCGA cancer types in order to identify the hcSLs and lcSLs (Methods), therefore the correlation result for all cSLs is also different from that in (a).

Similar to our earlier analysis, we see that the TCLs computed from the hcSLs correlate much stronger with onset age than those from the lcSLs or all cSLs (Spearman’s ρ = 0.603 vs −0.157, **Fig. 3b**, Fig. S7a), and also stronger than those obtained from control tests performed as before (empirical P < 0.001, Fig. S7b-d). As with the case of cancer risk, the observed correlation is dominated by the SL genetic interaction effects rather than the single gene effects (Fig. S7e-g, Supp. Note).

### The *activity state of cSL partners of some tumor suppressor genes predicts the specific tissues in which they are known to drive cancer*

Further investigating the role of cSLs in cancer development, we turned to ask whether cSL may also contribute to the tissue/cancer-type specificity of tumor suppressor genes (TSGs) (10,25). Specifically, we reasoned that loss-of-function mutations of a TSG during carcinogenesis will be less frequent in tissues where its *cSL partner genes* are lowly expressed, due to the synthetic lethal effect of such co-inactivation on the emerging cancer cells. To study this hypothesis, we obtained a list of TSGs together with the tissues in which their loss is annotated to have a tumor-driving function from the COSMIC database (11) (Table S6a). We further identified the cSL partner genes of each such TSG using ISLE (22) (Methods, Table S6b). In total, there are 23 TSGs for which we were able to identify more than one cSL partner gene. Consistent with our hypothesis, we find that in the majority of the cases, the cSL partner genes of TSGs have higher expression levels in the tissues where the TSGs are known drivers compared to the tissues where they are not established drivers (binomial test for the direction of the effect P = 0.023, **Fig. 4a**). We identified 10 TSGs whose individual effects are significant (FDR < 0.05) as well as cSL-specific (as shown by the random control test), and all these 10 cases exhibit the expected direction of effect (labelled in **Fig. 4a**, Table S6c; two example TSGs, *FAS* and *BRCA1*, are shown in **Fig. 4b**, details in Fig. S8, Methods). Reassuringly, these findings disappear under randomized control tests involving random partner genes of the TSGs and shuffled TSG-tissue type mappings (Supp. Note), further consolidating the role of cancer-specific cSLs of normal tissues in cancer risk and development.

**Figure 4.**
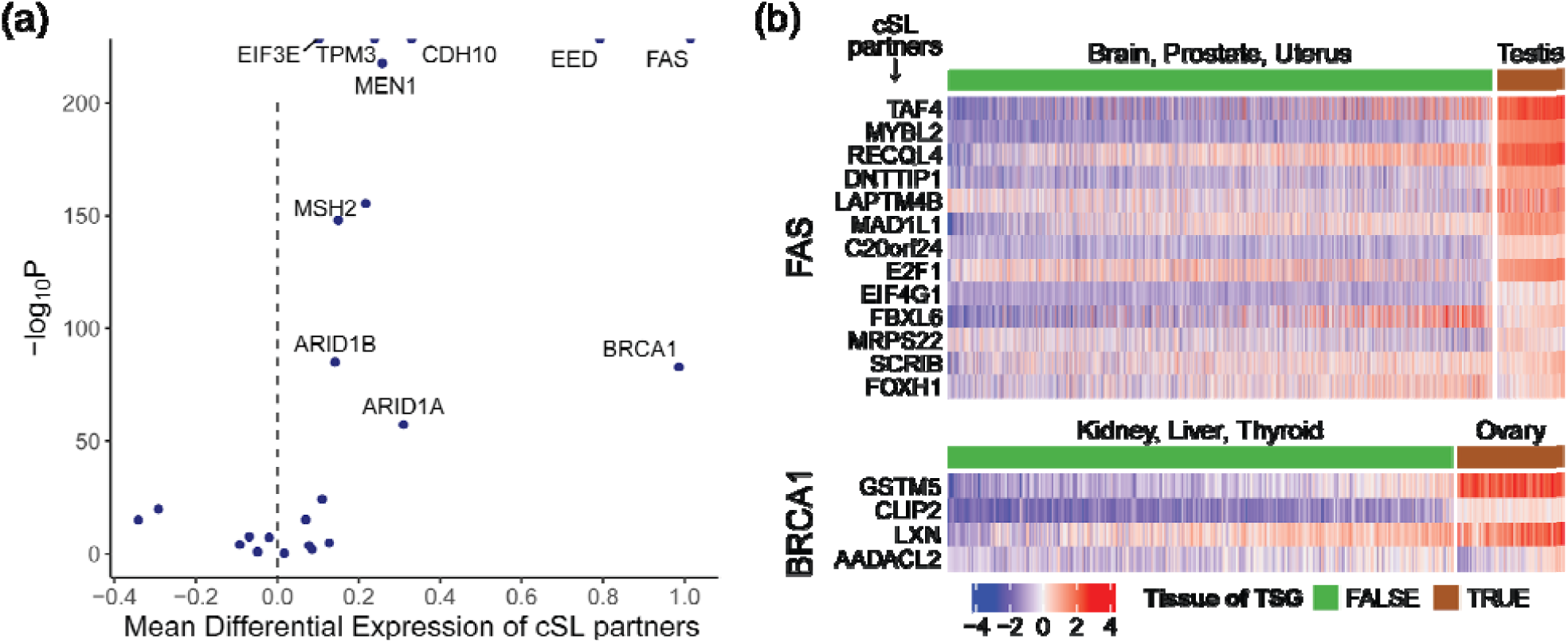
**(a)** For each tissue-specific tumor suppressor (TSG) gene G_i_, the expression levels of its cSL partner genes in the tissue type(s) where gene G_i_ is a TSG were compared to those where gene G_i_ is not an established TSG, using GTEx normal tissue expression data. The volcano plot summarizes the result of comparison with linear models. Positive linear model coefficients (X-axis) mean that the expression levels of the cSL partner genes are on average higher in the tissue(s) where gene G_i_ is a TSG. Many cases have near-zero P values and are represented by points (half-dots) on the top border line of the plot. Overall there is a dominant effect of the cSL partner genes of TSGs having higher expression levels in the tissues where the TSGs are known drivers (binomial test P = 0.023). All TSGs with FDR < 0.05 that also passed the random control tests are labeled. **(b)** Examples of two well-known TSGs, FAS and BRCA1, are given. The heatmaps display the normalized expression levels of their cSL partner genes (rows) in tissues of where these two genes are known to be TSGs (according to the annotation from the COSMIC database (11)) and in tissues where they are not established TSGs (columns), respectively. High and low expressions are represented by red and blue, respectively. For clarity, one typical tissue type where the TSG is a known driver (e.g. testis for FAS) and three other tissue types where the TSG is not an established driver (and the least frequently mutated) are shown.

## Discussion

In this work we show that the cSL load in normal tissues is a strong predictor of tissue-specific lifetime cancer risk, and is much stronger than the pertaining predictive power observed on the individual gene level. Consistently, we find that higher cSL load in the normal tissues from young people is associated with later onset of the cancers of that tissue. As far as we know, no other factor has been previously reported to be predictive of cancer onset age across tissues. Finally, we show that the activity status of cSL partners of tumor suppressor genes can explain their tissue-specific inactivation.

We have shown that the effect of cSL on the cancer risk (and cancer development in general) in normal tissues can be attributed to the fact that many of the cSLs are specific to cancer and have weak or no functional lethal effect in the normal tissues (Figs. 2d, 3b, Fig. S5), therefore normal tissues can bear relatively high cSL loads without being detrimentally affected -- quite to the contrary, they become more resistant to cancer due to the latent effect of these cSLs on potentially emerging cancer cells. Importantly, we emphasize that while we quantified the cSL loads using the normal tissue data from GTEx, the set of cSLs we used were derived exclusively in cancer from completely independent cancer datasets (and without using any information regarding lifetime cancer risk, onset, or tumor suppressor tissue specificity), so there is no circularity involved. The cSL load in normal tissues was computed to reflect the summed effects of individual cSL gene pairs. The underlying assumption is that the low expression of each cSL gene pair is synthetic sick (i.e. reducing cell fitness to some extent), and that the effects from different cSL gene pairs are additive, consistent with the ISLE method of cSL identification (22). Indeed, many experimental screenings of SL interactions also rely on techniques such as RNA interference that inhibits gene expression rather than completely knocks out a gene (26), and it is evident that most of the resulting SL gene pairs have milder than lethal effects. The additive cancer-specific lethal effect of such cSL gene pairs, however, can form a negative force impeding cancer development from normal tissues.

Obviously, as we are studying the across-tissue association between cSL load and cancer risk, it is essential to focus on cSLs that are common to many cancer types (i.e. pan-cancer). Therefore, we focused on cSLs identified computationally by ISLE via the analysis of the pan-cancer TCGA patient data (22). In contrast, most experimentally identified cSLs are obtained in specific cancer cell lines and are thus less likely to be pan-cancer (and possibly, less clinically relevant (22)). However, for completeness, we also compiled a set of experimentally identified cSLs from published studies (22,27) (Supp. Note, Table S1b). The corresponding TCL values computed using this set of cSLs correlates significantly with lifetime cancer risk, but not with cancer onset age; the correlation with cancer risk is also markedly weaker than that obtained from ISLE-derived cSLs (Spearman’s ρ = −0.433, P = 0.024, Fig. S9a, control tests and detailed analysis explained in Supp. Note). These experimentally identified cSLs can explain some cases of tissue-specific TSGs including BRCA1 and BRCA2 (Fig. S9e), but do not result in overall significant accountability for a large proportion of TSGs present in the analysis (like in Fig. 4a). This corroborates the importance of pan-cancer cSLs and their relevance to cancer risk.

Interestingly, tissue cSL load is not likely a corollary of the number of tissue stem cell divisions (NSCD) and DNA methylation (LADM; the latter was thought to be closely related to NSCD (6)), as cSL load is computed by analyzing bulk tissues, where stem cells occupy only a minor proportion. We have shown that TCL significantly adds to either NSCD or LADM in predicting lifetime cancer risk (Figs. 2b,c), which also suggests that cSL load is an independent factor of cancer risk acting via unique mechanisms. Furthermore, NSCD is measured as the product of the rate of tissue stem cell division and the number of stem cells residing in a tissue (2), and we confirmed that TCL is correlated with lifetime cancer risk independent of both of these components (partial Spearman’s rho −0.510 and −0.567, P = 0.007 and 0.002, respectively, Fig. S10a,b). We additionally tested that proliferation indices computed for the bulk normal tissues do not correlate with lifetime cancer risk across tissues (Spearman’s ρ = 0.062, P = 0.77, Fig. S10c, Supp. Note). Further, we verified that our observed correlations are not confounded by the number of samples from each cancer or tissue type (Fig. S11).

Taken together, our findings demonstrate the contribution of synthetic lethality to cancer risk, onset time, and context-specificity of tumor suppressors across human tissues. Beyond the effect on cancer after it has developed, our work highlights the role of cancer synthetic lethality during the entire course of carcinogenesis. While synthetic lethality has been attracting tremendous attention as a way to identify cancer vulnerabilities and target them, this is the first time that its role in mediating cancer development is uncovered.

## Methods

### Cancer SL (cSL) interaction networks

The cSL gene pairs computationally identified by the ISLE (identification of clinically relevant synthetic lethality) pipeline was obtained from (22). We used the cSL network identified with FDR < 0.2 for the main text results, containing 21534 cSL gene pairs, which is a reasonable size representing only about 1 cSL partner per gene on average. This also allows us to capture the effects of many weak genetic interactions. Nevertheless, we also used the cSL network with FDR < 0.1 (only 2326 cSLs) to demonstrate the robustness of the results to this parameter (Supp. Note). Each gene pair is assigned a significance score (the “SL-pair score” defined in Lee *et al*. 2018 (22)), that a higher score indicates that there is stronger evidence that the gene pair is SL in cancer. Out of these, we used 20171 cSL gene pairs whose genes are present in the GTEx data (Table S1a). The experimentally identified cSL gene pairs were collected from 18 studies (references in Supp. Note. Obtained from the Supplementary Data 1 of Lee *et al*. 2018 except for those from Horlbeck *et al*. 2018 (27)). Horlbeck *et al*. provided a gene interaction (GI) score for each gene pair in two leukemia cell lines. Gene pairs with GI scores < −1 in either cell line were selected as cSLs. A total of 27975 experimentally identified cSLs were obtained, out of which 27538 have both their genes present in the GTEx data (Table S1b).

### GTEx and TCGA data

The V6 release of Genotype-Tissue Expression (GTEx) (20) RNA-seq data (gene-level RPKM values) were obtained from the GTEx Portal (https://gtexportal.org/home/). The associated sample phenotypic data were downloaded from dbGaP (28) (accession number phs000424.vN.pN). For comparing the level of negative selection to co-inactivation of cSL gene pairs between normal and cancer tissues, the RNA-seq data of the Cancer Genome Atlas (TCGA) and GTEx as RSEM values that have been processed together with a consistent pipeline that helps to remove batch effects were downloaded from UCSC Xena (29). The expression data for each tissue type (normal or cancer) was normalized separately (inverse normal transformation across samples and genes) before being used for the downstream analyses. We mapped the GTEx tissue types to the corresponding TCGA cancer types (Table S2b), resulting in one-on-many mappings, e.g. the normal lung tissue was mapped to both lung adenocarcinoma (LUAD) and lung squamous cell carcinoma (LUSC).

### Cancer risk data and onset age

Lifetime cancer risk denotes the chance a person has of being diagnosed with cancer during his or her lifetime. Lifetime cancer risk data (Table S2a) are from Tomasetti and Vogelstein, 2015 (2), which are based on the US statistics from the SEER database (1). We derived the cancer onset age based on the age-specific cancer incidence data from the SEER database with the standard formula (30). Specifically, for each cancer type, SEER provides the incidence rates for 5-year age intervals from birth to 85+ years old. The cumulative incidence (CI) for a specific age range *S* is computed from the corresponding age-specific incidence rates (*IR*_*i*_, *i* ∈ *S*) as *CI* = 5Σ_*i*∈*S*_ *IR*_*i*_, and the corresponding risk is computed as *risk* = 1 - exp(-*CI*). The onset age for each cancer type (Table S3) was computed as the age when the CI from birth is 50% of the lifetime CI (i.e. birth to 85+ years old). Usually, the onset age defined as such is between two ages where the actual CI data is available, so the exact onset age was obtained by linear interpolation. Alternative parameters were used to define onset age (Supp. Note) in order to show the robustness of the correlation between tissue cSL load and cancer onset age based on different definitions.

### Computing cSL load

For each sample, we computed the number of cancer-derived SL gene pairs that have both genes lowly expressed, and divided it by the total number of cSLs available to get the cSL load per sample. In the ISLE method described by (22), low expression was defined as having expression levels below the 33 percentile in each tissue. Thus the ISLE-derived cSL gene pairs were shown to exhibit synthetic sickness effect when both genes in the gene pair are expressed at levels below the 33 percentile in each tissue, even though this appears to be a very tolerant cutoff (22). We therefore adopted the same criterion for low expression for the main results, although we also explored other low expression cutoffs to demonstrate the robustness of the results (Supp. Note).

### Computing tissue cSL load and correlation with lifetime cancer risk

Tissue cSL load (TCL) of each tissue type is the median value of the cSL loads of all the samples (or a subpopulation of samples) in that tissue, with cSL load of a sample computed as above. For example, TCL for the older population (age ≥ 50 years) is the median cSL load for the samples of age ≥ 50 years in each tissue type. For analyzing the correlation between the TCLs computed from GTEx normal tissues and cancer risk, we mapped the GTEx tissue types to the corresponding cancer types for which lifetime risk data are available from Tomasetti and Vogelstein, 2015 (2), resulting in 16 GTEx types mapped to 27 cancer types (Table S2a). Gallbladder non papillary adenocarcinoma, and Osteosarcoma of arms, head, legs and pelvis are not mapped to GTEx tissues and excluded from our analysis.

### Computing cSL single-gene load

To investigate the effect on the single gene level, we computed the cSL single-gene load in a paralleling way to the computation of the cSL load. Among all the unique genes constituting the cSL network (i.e. cSL genes), we computed the fraction of lowly expressed cSL genes for each sample as the cSL single-gene load, where low expression was defined in the same way as the computation of cSL load as elaborated above. Similarly, tissue cSL single-gene load is the median value of the cSL single-gene loads of all the samples in a tissue.

### Predicting tissue lifetime cancer risk with linear models

The lifetime cancer risks across tissue types were predicted with linear models containing three different sets of explanatory variables: (i) the number of total stem cell divisions (NSCD) alone, (ii) tissue cSL load alone, and (iii) NSCD together with tissue cSL load. Log-likelihood ratio (LLR) test was used to determine whether model (iii) (the full model) is significantly better than model (i) (the null model) in predicting lifetime cancer risks. The three models were also used to predict the lifetime cancer risks with a leave-one-out cross-validation procedure, and the prediction performances were measured by Spearman correlation coefficient. A similar analysis was performed to predict lifetime cancer risks across tissue types with three linear models involving the level of abnormal DNA methylation levels of the tissues (6): (i) the number of levels of abnormal DNA methylation (LADM) alone, (ii) tissue cSL load alone, and (iii) LADM together with tissue cSL load.

### Identifying and analyzing highly specific and lowly specific cSLs (hcSLs and lcSLs)

For each pair of GTEx normal-TCGA cancer of the same tissue type (Table S2b), we computed the fraction of samples where a cSL gene pair *i* has both genes lowly expressed (defined above) among the normal samples (*fn*_*i*_) and cancer samples (*fc*_*i*_), and computed a *specific score* as *rs*_*i*_ = *fn*_*i*_ *-fc*_*i*_. We selected the hcSLs as those whose specific scores are greater than the 75% percentile of all scores, and lcSLs as those with a score below the 25% percentile (Table S4a,b). We compared SL significance scores between the hcSLs and lcSLs in each tissue using a Wilcoxon rank-sum test. For each type of the GTEx normal tissues used in this analysis (i.e. those that can be mapped to TCGA cancer types), we also computed the tissue cSL load as above but using the hcSLs, lcSLs, or all cSLs, respectively, and analyzed their correlation with lifetime cancer risk or cancer onset age across the tissues.

### Pathway enrichment of the hcSLs

We designed an empirical enrichment test as below to account for the fact that each cSL consists of two genes. For the hcSLs in each tissue type and each given pathway from the Reactome database (31), we computed the odds ratio (OR) for the overlap between the genes in hcSLs and the genes within the pathway based on the Fisher’s exact test procedure, with the “background” being all the genes in the ISLE-inferred cSLs. A greater than 1 OR indicates that the hcSLs are positively enriched for the genes of the pathway. To determine the significance of the enrichment, we repeatedly and randomly sampled the same number of cSLs as that of the hcSLs, computed the ORs similarly, and computed the empirical P value as the fraction of cases where the OR from the random cSLs is greater than that from the hcSLs. We corrected for multiple testing across pathways with the Benjamini-Hochberg method.

### Analyzing the tissue-specificity of tumor suppressor genes

We obtained the list of TSGs and their associated tissue types from the COSMIC database (11) (https://cancer.sanger.ac.uk/cosmic/download, the “Cancer Gene Census” data. Table S6a). For each TSG, their cSL partner genes were identified using the ISLE pipeline (22) with an FDR cutoff of 0.1 (Table S6b). Here the FDR cutoff is more stringent than that used for the pan-cancer genome-wide cSL network (FDR < 0.2 for the main results) since here FDR correction was performed for each TSG, corresponding to a much lower number of multiple hypotheses. As a result, the FDR correction has more power and a relatively more stringent cutoff can give rise to a more reasonable number of cSL partner genes per TSG. We focused our analysis on 23 TSGs for which more than one cSL partner genes were identified (no cSL partner was identified for most of the other TSGs). The expression levels of the cSL partner genes were then compared between tissue type(s) where the TSG is a known driver and the rest of the tissues where the TSG is not an established driver with linear models. Specifically, the expression levels of the cSL partners were modeled with two explanatory variables: (i) driver status of the TSG in the tissue (binary) and (ii) cSL partner gene (categorical, indicating each of the cSL partner genes of a TSG). The coefficient and P value associated with variable (i) were used to analyze the general trend of differential expression among the cSL partner genes. Positive coefficients of variable (i) means that the expression levels of the cSL partner genes are on average higher in the tissue(s) where the TSG is a known driver compared to those in the tissues where the TSG is not an established cancer driver.

## Data/Code availability

The R code and the relevant data are available at https://hpc.nih.gov/~chengk6/SL_cancer_risk.zip.

## Acknowledgments

This research was supported in part by the Intramural Research Program of the National Institutes of Health (NIH), National Cancer Institute and the Center for Cancer Research. This work utilized the computational resources of the NIH HPC Biowulf cluster (http://hpc.nih.gov). The Genotype-Tissue Expression (GTEx) Project was supported by the Common Fund of the Office of the Director of the National Institutes of Health, and by NCI, NHGRI, NHLBI, NIDA, NIMH, and NINDS. We thank Dr. Stephen J. Chanock, Dr. Brid M. Ryan, Dr. Tom Misteli, Prof. Sridhar Hannenhalli, Dr. Xin Wei Wang, Dr. Curtis C. Harris, Dr. Alejandro Schaffer, Dr. Michael Gertz, Dr. Kun Wang, Sushant Patkar, Sanju Sinha, and Dr. Fiorella Schischlik for their critical comments and suggestions on our study and the manuscript. K.C. is supported by the NCI-UMD Partnership for Integrative Cancer Research fellowship.

## Author contributions

KC, NUN, ER formulated the research question and study design. KC, NUN carried out the analysis. JSL provided help with the computational cSL prediction pipeline. KC, NUN, ER analyzed the results. KC, NUN, ER wrote the manuscript with inputs from JSL. ER supervised the research. The manuscript has been read and approved by all the authors.

## Author Information

Eytan Ruppin is a co-founder and scientific consultant of Pangea Therapeutics (https://pangeamedicine.com/) which focuses on precision oncology and synthetic lethality; however he has divested all his shares and receives no salary or financial benefit from this company. The work in this manuscript is not related to the work of this company. The other authors declare no conflict of interest. Correspondence and requests for materials should be addressed to.

