## Supplementary notes and figures for "Synthetic lethality across normal tissues is strongly associated with cancer risk, onset, and tumor suppressor specificity"

#### Details of the cSL gene pairs

The cSLs obtained from Lee *et al.*[1] were computationally identified with the ISLE (identification of clinically relevant synthetic lethality) pipeline, which mined data from 8749 cancer TCGA patient tumors across 28 cancer types and applied a 4 step procedure to identify the cSL network. The pipeline starts with an initial set of experimentally identified cSL interactions either by double knock-out/knock-down in a single cell line or by inference from the single RNAi/CRISPR screens in a large number of cancer cell lines. Then, it analyzes patient tumor data across the 28 cancer types to further filter those pairs that show the evidence of (i) negative selection of and (ii) better patient survival due to co-inactivation while controlling for the effect of individual genes, followed by (iii) a phylogenetic screen (described in detail in Lee *et al.*[1]). We used the cSL network identified with FDR < 0.2 and FDR < 0.1 as described in the Methods. Experimentally identified cSLs were obtained by pooling the experimentally identified cSL pairs from 17 double knock-down/knock-out screens[2-18] and combining it with the cSL pairs identified with CRISPRi experiments by Horlbeck *et al.*[19] as described in the Methods. The merit of using ISLE-inferred cSL pairs over the experimentally identified cSLs is that the selected cSL pairs may be more likely to be clinically relevant across many cancer types and less likely to be cancer-type specific.

**Effect size measure of Wilcoxon tests and multiple testing**

Throughout the text we measured the effect size of the Wilcoxon tests by the rank-biserial correlation, which ranges from -1 and 1. Strong effects are represented by rank-biserial correlations whose absolute values are close to 1, and weak effects represented by values close to 0. The exact formula for calculating the rank-biserial correlation can be found in Wendt *et al.*[20] We corrected for multiple testing using the Benjamini-Hochberg method throughout our study.

**Robustness of the correlation between tissue cSL load and lifetime cancer risks, and cancer onset age**

*Alternative cSL load measures*

We used various alternative approaches to compute cSL load and correlated them with lifetime cancer risk in order to show the robustness of our findings. These alternative approaches are described below:

a) We computed the correlation between lifetime cancer risk and cSL load (for older populations) using a more stringent FDR threshold (FDR < 0.1 instead of FDR < 0.2) for selecting the cSL network using ISLE (leading to 2326 pairs). We still get a significant correlation (Spearman's rho = -0.613, P = 0.000668, Fig. S1a).
b) We removed genes that have zero expression in more than 90% of all GTEx samples. We then computed cSL load (for older populations) with the remaining genes, and found a significant correlation with lifetime cancer risk (Spearman's rho = -0.605, P = 0.000838, Fig. S1b). Here the ISLE-inferred cSLs with FDR < 0.2 were used.
c) For our main results presented in Fig. 2 of the main text, tissue cSL load (TCL) was computed by taking the median (i.e. the 50 percentile) cSL load of the samples

from a tissue. We additionally computed TCL by taking the 65 percentile or the 40 percentile of the cSL loads in the samples (age ≥ 50 years) of a tissue. TCLs obtained as such are also significantly correlated with lifetime cancer risk (rho = -0.794, P = 7.59e-7, Fig. S1c; and rho = -0.52, P = 0.0054, Fig. S1d, respectively).

d) We have defined a cSL gene pair to be inactivated if the expression levels of both genes in the cSL pair are below the 33 percentile within the population (i.e. each tissue type). We also used definition of inactivated cSLs based on other percentiles, including 40 percentile and 20 percentile. The corresponding correlations with lifetime cancer risk are shown in Fig. S1e and S1f, respectively, which remain significant.

Taken together, these results show that tissue cSL load is robustly (negatively) correlated with lifetime cancer risk across tissues.

*Alternative mappings between normal and cancer tissue types*

In the main result as presented in Fig. 2 of the main text, we used the mapping between 16 GTEx normal tissues to 27 cancer types as given in Table S2a. However certain mappings were not exact due to the limitations of the data available, for example, salivary gland was the closest GTEx tissue type that may be mapped to head and neck carcinoma (with or without HPV-16), but in a very non-strict sense. Even if we removed these two mappings, we still got a similar significant correlation between tissue cSL load (TCL) with cancer lifetime risk (Spearman’s rho = -0.662, P = 0.00031, Fig. S1g). As another example, we additionally removed the mapping between “Brain - Cerebellum” in GTEx and Medulloblastoma, and still get a good correlation between TCL and risk (Spearman’s rho = -0.623, P = 0.00067, Fig. S1h).

*Alternative cancer onset age definition*

We computed the onset age for each cancer type from SEER incidence data[21]. We mapped the cancer types in SEER to the tissue type in GTEx, given in Table S3. We then computed TCL as described above for the samples below 40 years old in each GTEx tissue, and correlated the TCL with cancer onset age across tissues (for the result in Fig. 3 of main text). TCLs were also computed for other upper cutoffs of ages (including 45 and 50 years old), and their correlations with onset age are shown in Fig. S6, in order to show that the result is robust to the parameters used. We find that in general, TCL for younger populations correlates positively with cancer onset age in a robust way. This is consistent with the hypothesis that the lower the TCL for a tissue, the earlier the cancer onset time, thus showing that tissue cSL load can impede cancer development.

**Randomized control analysis**

We did three types of random control analysis to test that the various observed effects in this study are specific to the cSL gene pairs and are not random. The three random controls were performed using: (i) random gene pairs; (ii) shuffled cSL gene pairs; and (iii) degree-preserving randomized cSL network (same size as the actual cSL network). Below we describe the detailed procedure testing for the cSL load correlation with cancer risk as an example.

(i) Random gene pairs: we randomly sampled 20171 gene pairs from all the genes in the GTEx data and computed a tissue "pseudo-cSL load" with these random gene pairs, for each tissue for the older population (≥ 50 years), and then computed its correlation with lifetime cancer risk. This procedure was repeated for 1000 iterations and a histogram showing correlation distributions are shown in Fig. S3a. We see that the correlations are not significant (median Spearman's rho = -0.21, P = 0.29). An empirical p-value was computed using the correlation obtained from the actual cSL network and the correlation distribution of the tissue pseudo-cSL loads (P < 0.001, Fig. S3a), indicating that the signal from the true cSL network is always much stronger than random signals.

(ii) Shuffled cSL gene pairs: starting from the 20171 cSL gene pairs listed in a table (20171-by-2, each row is a gene pair), we randomly shuffled the columns of the table to obtain a list of shuffled gene pairs, and computed a tissue pseudo-cSL load for each GTEx tissue for older populations (age ≥ 50 years). This was correlated with cancer lifetime risk. This procedure was repeated for 1000 iterations and a histogram showing correlation distributions are shown in Fig. S3b. We see that many of these random correlations are significant (median Spearman's rho = -0.537, P = 0.0039). This can be due to an effect on the single gene level, which we described in the main text, and showed that the genetic interaction effect actually is the dominant one (vs the single-gene effect,

Fig. S3d-g). For more details on the single-gene effect analysis, see the section "*Details on the single-gene effect analysis*" below. Additionally, the empirical p-value for the randomization test computed as above is P < 0.001 (Fig. S3b), indicating that the signal from the true cSL network is still the strongest.

(iii) Degree-preserving randomized cSL network: starting from the actual cSL network consisting of 20171 cSL gene pairs (edges), we performed a degree-preserving randomization, rewiring the network but retaining the original degree distribution. Tissue pseudo-cSL load was computed as above and correlated with cancer lifetime risk. This procedure was repeated for 1000 iterations and a histogram showing correlation distributions are shown in Fig. S3c. The random correlations have a median value of Spearman's rho = -0.537, P = 0.0039, which as elaborated above and further explored in the main text, can be due to the single-gene effect (Fig. S3d-g). Nevertheless, again the empirical p-value is P < 0.001 (Fig. S3c), indicating a strong and robust cSL-specific effect.

Similarly, such control tests were performed for the correlation with cancer onset age (Fig. S7) and experimentally identified cSLs (Fig. S9). We also performed control tests for the analysis of the tissue type-specificity of tumor suppressor genes (TSGs). As shown in Fig. 4 of the main text, the true cSL partner genes of a TSG tend to have higher expression in the tissue type(s) where the TSG is a known driver and the rest of the tissue types where the TSG is not an established cancer driver, supporting our hypothesis that the tissue type-specificity of TSGs can be (partly) explained by cSL. Specifically, out of the 23 cases of TSGs, 17 of them show such a trend, and only 6 cases show the opposite trend but with much weaker effect sizes and P values. We simply uses the difference in the number of cases on both sides as a test statistics (with the true cSL pairs, the value is 17-6=11), with a larger value of the statistics indicating a stronger evidence supporting our hypothesis. For each TSG, given its true cSL partner genes identified by the ISLE method, we randomly sampled the same number of genes from all the genes as its "pseudo-partner genes", and we analyzed the differential expression (DE) of these pseudo-partner genes between the tissue type(s) where the TSG is a known driver and the rest of the tissue types where the TSG is not an established cancer driver using linear model, as described in Methods. Summarizing the results of all TSGs, the test statistics was calculated. After repeating the random sampling, we computed the empirical P value based on the fraction of times that the statistics obtained using pseudo-partner genes is greater than that obtained using the true cSL partners. We also performed another control test by randomly shuffling the mapping between the TSG and the tissue types where they have established driver functions, and the empirical P value was computed in a similar fashion. Both control tests yielded empirical $P < 0.05$. Additionally, we also performed the control tests as above for each of the TSG, with the test statistics being the linear model coefficient, which represents the mean difference in the expression level of the cSL partner genes between the tissue(s) of the TSG and the tissues where the TSG is not a known driver, and the empirical P values obtained for each TSG is given in Fig. S8.

**Details on the single-gene effect analysis**

As we described above and in the main text, some of the random control tests yielded significant signals in terms of, e.g. correlation with cancer risk or onset age, although the signals from the actual cSL gene pairs are always much stronger. Notably, it is the shuffled cSL gene pairs or the degree-preserving randomization that produced significant signals, but not the purely random gene pairs. We therefore reasoned that the genes within the cSL network (*cSL genes*) may have an effect on the single gene level, and although the cSL load is meant to capture the pairwise cSL genetic interaction effect, it actually also captures some of the single-gene effect due to the way it is computed. Accordingly we computed the *tissue cSL single gene load (SGL)* as described in the main text, and confirmed the single-gene effect wrt cancer risk (Fig. S3d) and onset age (Fig. S7e). Interestingly in both cases, the single-gene effect is only exhibited by the cSL genes, and no significant signal can be found using random sets of genes or all genes (Fig. S3e-f, S7f), which could be due to additional weak genetic interaction effects among the cSL genes not captured by the current cSL network. Further, as described in the main text, we examined the partial correlation between tissue cSL load (TCL) and risk/onset age after controlling for the SGL, and showed that it is the cSL genetic interaction effect that dominates the observed correlations (Fig. S3g, S7g). Specifically, we regressed out the SGL component from both TCL and cancer risk/onset age with linear regression, then correlated the residues. This is the standard procedure of partial correlation and is equivalent to a multiple regression model for cancer risk/onset (as the dependent variable) with both SGL and TCL as the independent variables (covariates). We adopted the partial correlation procedure here (instead of the multiple regression) since it gives the correlation value, which is easier to compare to the rest of our results.

**Reproducing the result of Tomasetti & Vogelstein[22] and Klutstein *et al.*[23]**

For our analysis, only a subset of the cancer types in Tomasetti & Vogelstein 2015[22] have matched normal tissue types available from the GTEx database (Table S2a), as explained in the Methods. To facilitate comparison, we reproduced the result of Tomasetti & Vogelstein 2015[22] that there is a strong association between the number of tissue stem cell divisions with cancer lifetime risk only on this subset of tissue types (Spearman's rho = 0.717 , P = 2.57e-5, Fig. S4). As a result of the drop-out of cancer types, this result is numerically different from that reported in their original manuscript[22] (Spearman's rho = 0.81; P = 3.5e-8). Similar situation exists for reproducing the result of Klutstein *et al.*[23] (Fig. S4).

**Computing the level of negative selection of cSL gene pairs**

For each gene, we define that it's lowly expressed in a sample if its expression level is below the 33 percentile of those in all samples of the same tissue type, which is consistent with the approach in Lee *et al.*[1] and as elaborated in the main text Methods. For each cSL gene pair, we computed the fraction of samples where both genes are lowly expressed for each of the normal and cancer tissue types, which were compared to those computed with random gene pairs with Wilcoxon rank-sum tests. Rank-biserial correlation, the effect size metric of Wilcoxon test was used to measure the level of negative selection, since effectively this metric will take zero values on average for random gene pairs, and negative values for gene pairs whose co-downregulation in a tissue is selected against. Also, a smaller negative value (i.e. larger absolute value) indicates a stronger negative selection.

**Details on the analysis of highly specific vs lowly specific cSLs (hcSLs and lcSLs)**

Note that for each tissue type, a distinct set of hcSLs and lcSLs were identified. The sets of hcSLs of the tissues have pairwise Jaccard indices ranging from 0.136 to

0.251 with a mean value of 0.163, which represents significantly more overlap compared to random sets of the same size ($P < 0.001$ in a randomization test for the mean Jaccard index). For the set of hcSLs and lcSLs for each tissue, we obtained the unique genes within the cSL pairs and identified the pathways they are enriched in as described in Methods. We focused on the pathways that are specifically enriched for the hcSL genes rather than the lcSL genes, since as we have shown the hcSLs are the ones most relevant in cancer risk.

**Contribution of cSL to the tissue-specificity of tumor suppressor genes (TSGs)**

A recent study reported that the expression levels of the genetic interaction (including cSL) partners of cancer driver genes differ across cancer types, in a way that is dependent on the tissue-specific functional status of the driver gene[24]. Their analysis was performed in cancer tissues, and thus does not directly support the role of cSL in cancer development. Our approach is different, however, in that we investigated the expression levels of the cSL partners of the TSGs in normal, non-cancerous tissues, and we sought to establish the link between the co-inactivation status of cSL gene pairs in normal tissues and the tissue type-specificity of carcinogenesis.

**The correlation between tissue cSL load and lifetime cancer risk is not confounded by the number (or rate) of normal tissue stem cell division**

Klutstein *et al.*[23] have suggested a link between abnormal DNA methylation and DNA replication in stem cells, consistent with the strong correlation they observed between LADM and NSCD. On the contrary, we showed in the main text that tissue cSL load (TCL) significantly adds to either NSCD or LADM in terms of the correlation with lifetime cancer risk and is likely not a corollary of tissue stem cell divisions. We further performed partial correlation of TCL and lifetime cancer risk, conditioned on either the rate of tissue stem cell division or the number of stem cells residing in each normal tissue (both obtained from Tomasetti *et al.*[22]) and found that strong correlations remain (Fig. S10a,b). We also computed the proliferation index (PI) for each of the samples in the GTEx normal tissue from gene expression using a proliferation signature named meta-PCNA[25]. PI was computed as the median expression values of the set of meta-PCNA genes defined in Venet *et al.* 2011[25], and is regarded as a proxy for the rate of cell proliferation within a sample. We then took the median value of the PI's for all samples of a tissue type, and correlated the median PI of a tissue with its lifetime cancer risk (across tissues), showing that there's no significant correlation (Fig. S10c). Also, we note that stem cell proliferation *rates* (rather than the total number of stem cell divisions), as estimated by Tomasetti *et al.*[22], correlates only weakly with lifetime cancer risk (Spearman's rho = 0.386, P = 0.046).

### Supplementary Tables

**Table S1**. ISLE-inferred and experimentally identified cSL gene pairs

**Table S2**. Tissue cSL load in the older population and its correlation with lifetime cancer risk

**Table S3**. Correlation between tissue cSL load in the younger population and cancer onset age

**Table S4**. Highly specific vs lowly specific cSLs_the TCLs computed from them and correlation with cancer risk and onset age

**Table S5**. Pathways specifically enriched by the highly specific cSLs in each tissue

**Table S6**. Tissue-specific tumor suppressor genes and their cSL partner genes

### Supplementary Figures

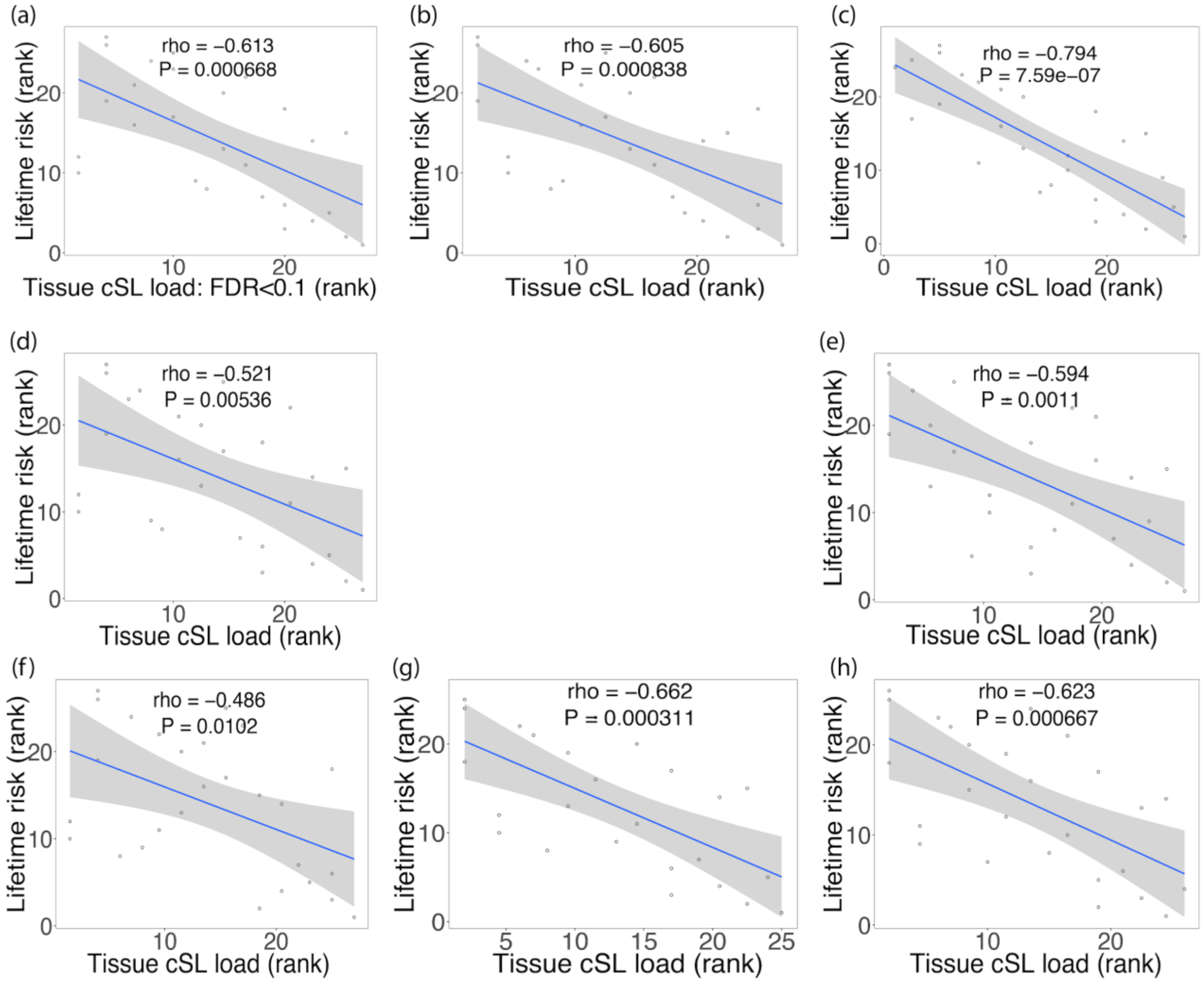


**Figure S1.** *Scatter plots showing Spearman correlations (rho) between cancer lifetime risk and: (a) cSL load (computed using FDR<0.1); (b) cSL load (computed by removing genes which have zero expression in more than 90% of the samples); (c) cSL load (by taking 65 percentile cSL load value across samples for each tissue, instead of median value); (d) cSL load (by taking 40 percentile cSL load value across samples for each tissue, instead of median value); (e) cSL load (using low expression threshold as less than 40 percentile); (f) cSL load (using low expression threshold as less than 20 percentile); (g) cSL load (removed 2 mappings between Head & neck squamous cell carcinoma and salivary gland); (h) cSL load (removed data point which mapped*

*Medulloblastoma to Brain-Cerebellum). cSL analysis was done on older populations for all sub-figures.*

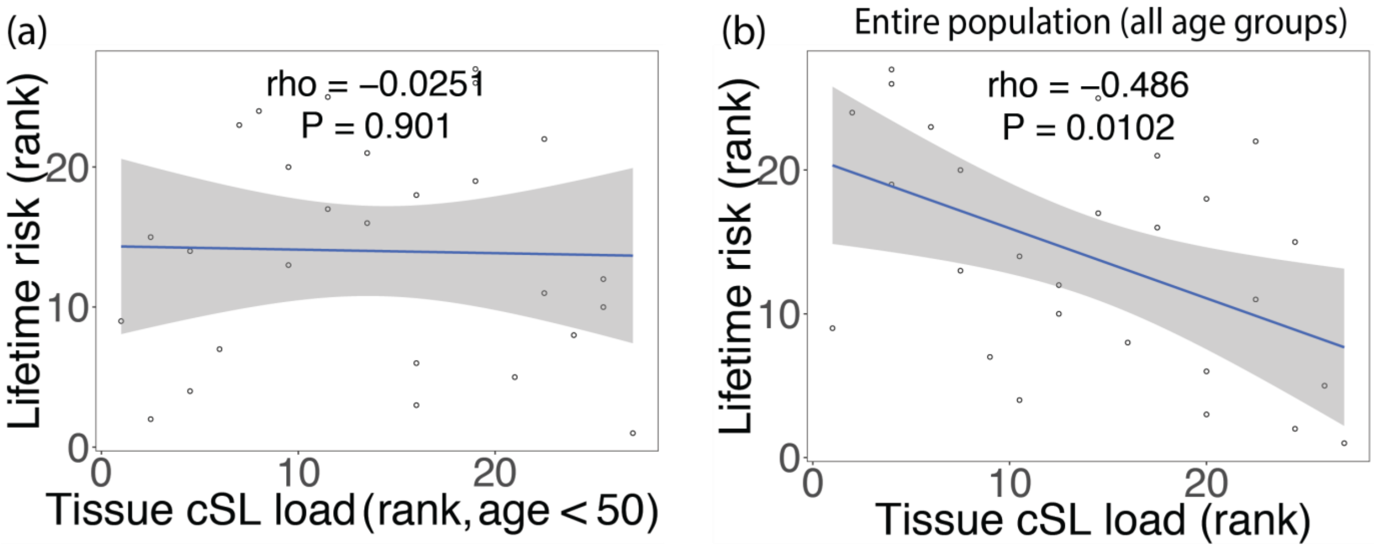


**Figure S2.** *Scatter plots showing Spearman correlations (rho) between cancer lifetime risk and: (a) Tissue cSL load (younger population, age < 50 years); (b) Tissue cSL load (on entire population which includes all age groups).*

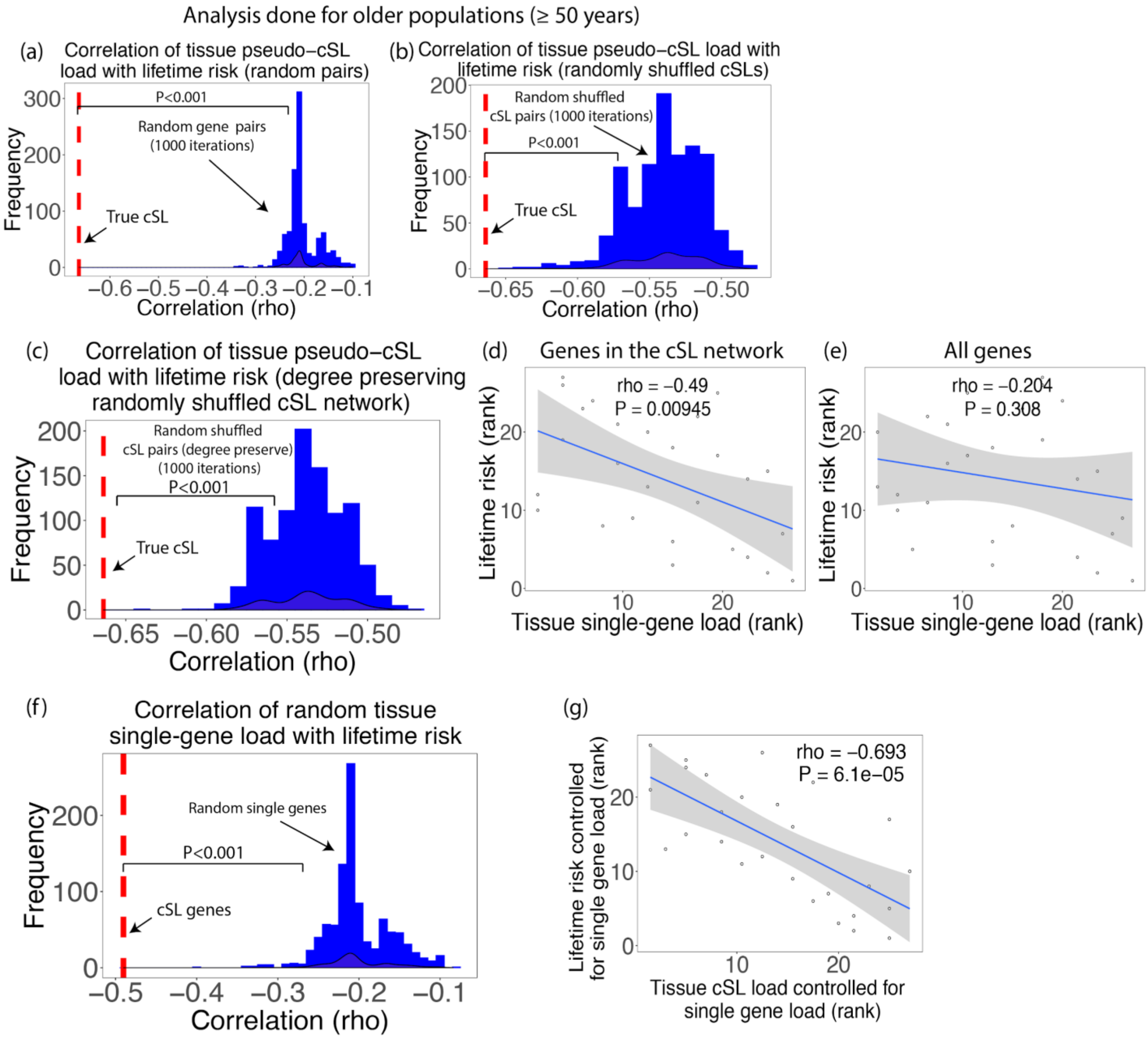


**Figure S3.** *(a) Histogram showing Spearman correlations of pseudo-cSL load (computed from random gene pairs) with cancer lifetime risk (no. of iterations = 1000). Correlation for tissue cSL load (True ISLE-inferred cSL) and cancer risk is shown in red. Randomization test shows that the correlation obtained from ISLE-inferred tissue cSL network is always more significant than those obtained from tissue pseudo-cSL loads (P < 0.001). (b) Histogram showing Spearman correlations of pseudo-cSL load (computed from shuffled cSL pairs) with cancer lifetime risk (no. of iterations = 1000). Correlation for*

*tissue cSL load (True ISLE-inferred cSL) and cancer risk is shown in red. Randomization test shows that the correlation obtained from ISLE-inferred cSL network is much more significant than those obtained from tissue pseudo-cSL loads (P < 0.001). (c) Histogram showing Spearman correlations of pseudo-cSL load (computed from degree-preserving randomized cSL network) with cancer lifetime risk (no. of iterations = 1000). Correlation for tissue cSL load (True ISLE-inferred cSL) and cancer risk is shown in red. Randomization test shows that the correlation obtained from ISLE-inferred cSL network is much more significant than those obtained from tissue pseudo-cSL loads (P < 0.001). Scatter plots showing Spearman correlations (rho) between cancer lifetime risk and: (d) tissue single-gene load (computed from only the genes in the ISLE-inferred cSL network); (e) tissue single-gene load (computed from all genes). (f) Histogram showing Spearman correlations of tissue single-gene load (computed from random single genes) with cancer lifetime risk (no. of iterations = 1000). Correlation for tissue single-gene load (inferred from ISLE-inferred cSL genes) and cancer risk is shown in red. Randomization test shows that the correlation obtained from cSL genes is much more significant than those obtained from random single genes (P < 0.001). (g) Scatter plots showing Spearman correlations (rho) between cancer lifetime risk and tissue cSL load controlled for tissue single-gene load (single-gene load is computed from only the genes in the ISLE-inferred cSL network).*

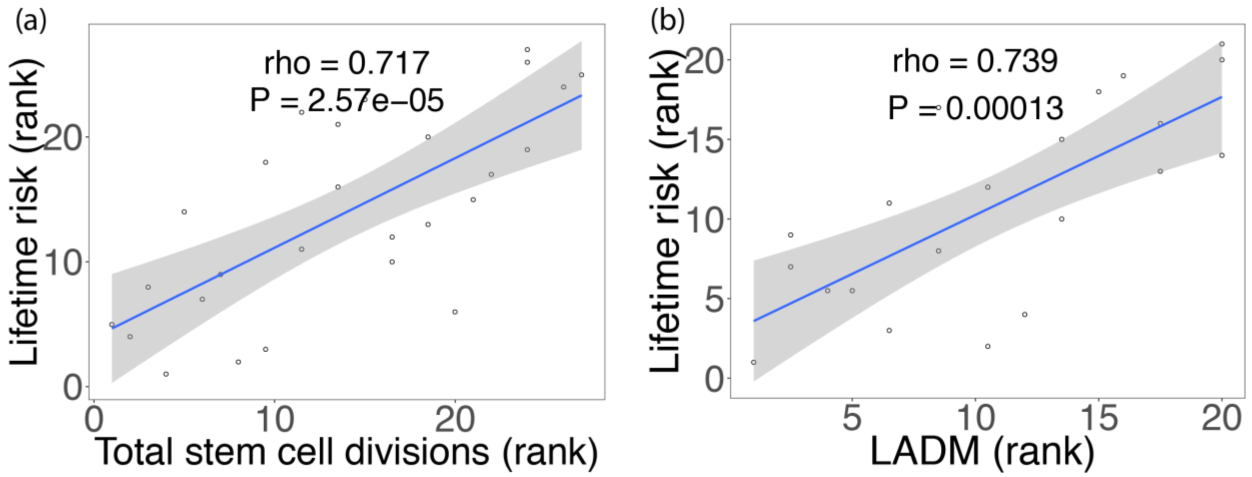


**Figure S4.** *Scatter plots showing Spearman correlations (rho) between lifetime cancer risk and: (a) total no. of stem cell divisions (NSCD); (b) LADM (levels of abnormal DNA methylation), across tissues.*

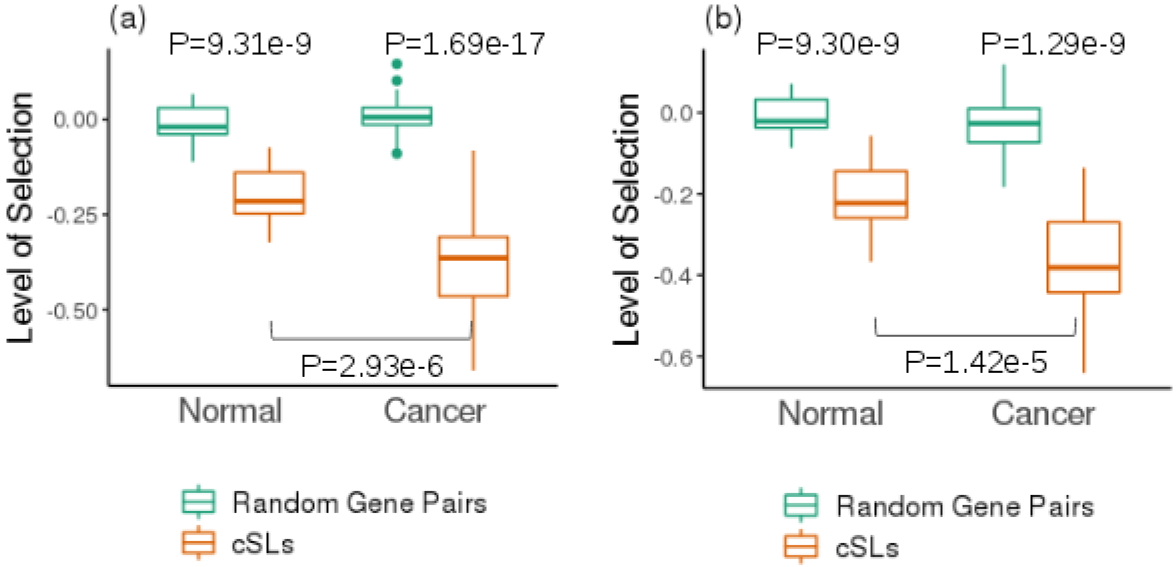


**Figure S5.** *(a) The level of negative selection (Supp. Note) against the inactivation of both genes in cancer-derived cSL gene pairs, compared to that of random gene pairs in both normal tissues from GTEx and cancers from TCGA. Co-inactivation of the genes in cSL gene pairs are much weaker selected against in GTEx normal tissues than in TCGA cancer samples. (b) Similar to (a), but with the analysis performed using cross-validation. ISLE was applied to a random subset of TCGA cancer samples to identify the cSLs, and the negative selection level in cancer was computed for the held-out samples not used by ISLE. P values for one-sided Wilcoxon rank-sum tests are shown.*

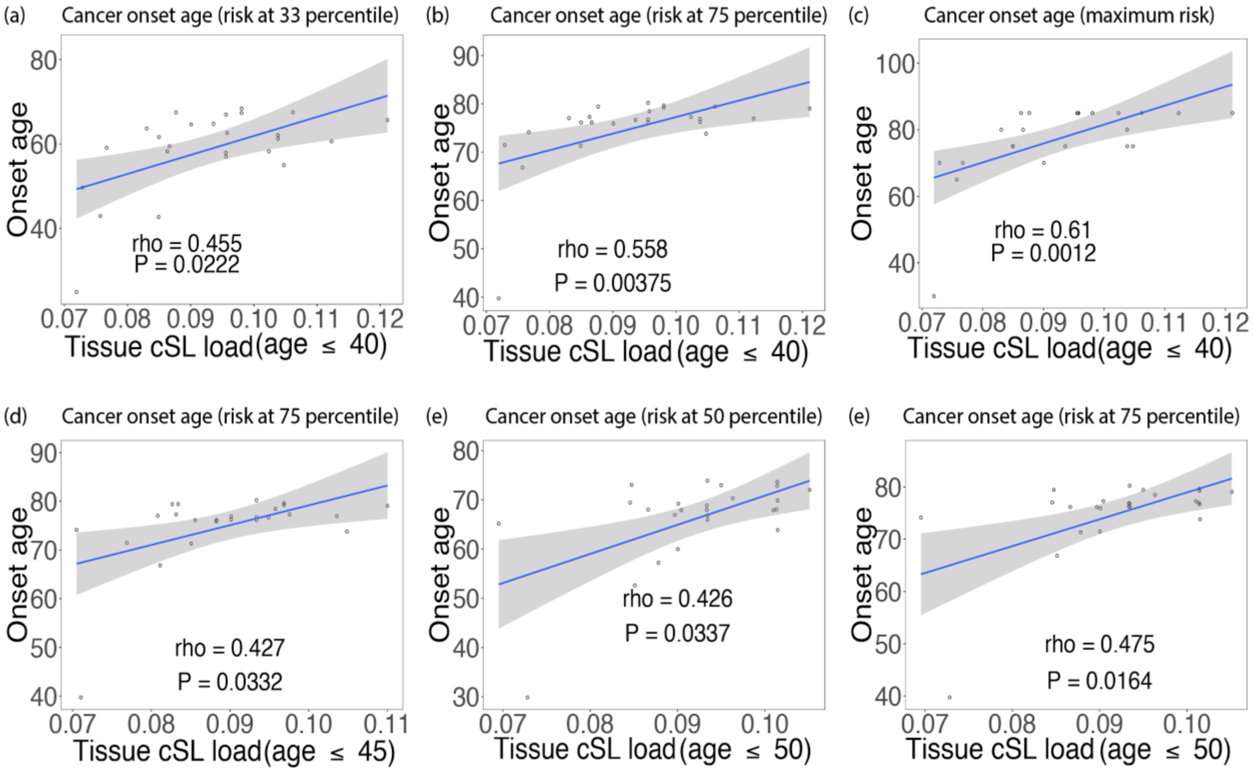


**Figure S6.** *Spearman correlation between Tissue cSL load and cancer onset age computed for different thresholds. Scatter plots showing correlation between: (a) Tissue cSL load (age ≤ 40) and cancer onset time (computed at 33 percentile of maximum risk); (b) Tissue cSL load (age ≤ 40) and cancer onset time (computed at 75 percentile of maximum risk); (c) Tissue cSL load (age ≤ 40) and cancer onset time (computed at maximum risk); (d) Tissue cSL load (age ≤ 45) and cancer onset time (computed at 75 percentile of maximum risk); (e) Tissue cSL load (age ≤ 50) and cancer onset time (computed at 50 percentile of maximum risk); (f) Tissue cSL load (age ≤ 50) and cancer onset time (computed at 75 percentile of maximum risk). All of them have FDR < 0.1.*

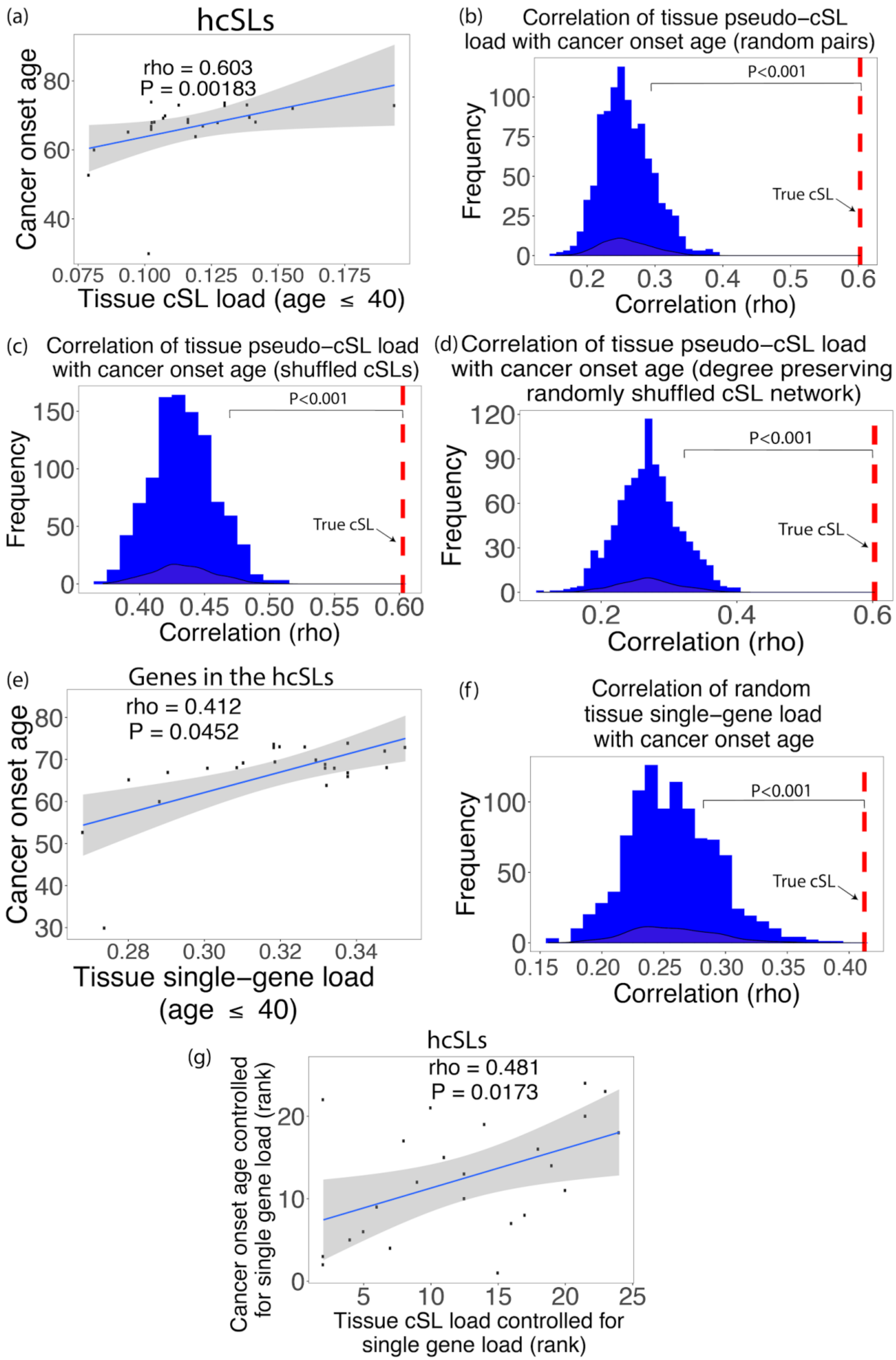
(a)
hcSLs
rho = 0.603
P = 0.00183
Cancer onset age
Tissue cSL load (age ≤ 40)
(b)
Correlation of tissue pseudo−cSL load with cancer onset age (random pairs)
P<0.001
True cSL
Frequency
Correlation (rho)
(c)
Correlation of tissue pseudo−cSL load with cancer onset age (shuffled cSLs)
P<0.001
True cSL
Frequency
Correlation (rho)
(d)
Correlation of tissue pseudo−cSL load with cancer onset age (degree preserving randomly shuffled cSL network)
P<0.001
True cSL
Frequency
Correlation (rho)
(e)
Genes in the hcSLs
rho = 0.412
P = 0.0452
Cancer onset age
Tissue single−gene load (age ≤ 40)
(f)
Correlation of random tissue single−gene load with cancer onset age
P<0.001
True cSL
Frequency
Correlation (rho)
(g)
hcSLs
rho = 0.481
P = 0.0173
Cancer onset age controlled for single gene load (rank)
Tissue cSL load controlled for single gene load (rank)

**Figure S7.** *(a) Scatter plot showing Spearman correlations (rho) between cancer onset age and tissue cSL load (computed from ISLE-inferred hcSL gene pairs). (b) Histogram showing Spearman correlations of pseudo-cSL load (computed from random gene pairs) with cancer onset age (no. of iterations = 1000). Correlation for tissue cSL load (True ISLE-inferred lcSL network) and cancer onset age is shown in red. Randomization test shows that the correlation obtained from ISLE-inferred hcSL gene pairs is always more significant than those obtained from pseudo-cSLs (P < 0.001). (c) Histogram showing Spearman correlations of pseudo-cSL load (computed from shuffled hcSL gene pairs) with cancer onset age (no. of iterations = 1000). Correlation for tissue cSL load (True ISLE-inferred hcSLs) and cancer onset age is shown in red. Randomization test shows that the correlation obtained from ISLE-inferred hcSL gene pairs is much more significant than those obtained from tissue pseudo-cSL loads (P < 0.001). (d) Histogram showing Spearman correlations of pseudo-cSL load (computed from degree-preserving randomized hcSL network) with cancer onset age (no. of iterations = 1000). Correlation for tissue cSL load (True ISLE-inferred lcSL) and cancer onset age is shown in red. Randomization test shows that the correlation obtained from ISLE-inferred hcSL network is much more significant than those obtained from tissue pseudo-cSL loads (P < 0.001). (e) Scatter plot showing Spearman correlations (rho) between cancer onset age and tissue single-gene load (computed from only the genes in the ISLE-inferred hcSL network). (f) Histogram showing Spearman correlations of tissue single-gene load (computed from random single genes) with cancer onset age (no. of iterations = 1000). Correlation for tissue single-gene load (inferred from ISLE-inferred hcSL genes) and cancer onset age is shown in red. Randomization test shows that the correlation obtained from hcSL genes is much more significant than those obtained from random single genes (P < 0.001). (g) Scatter plots showing Spearman correlations (rho) between cancer lifetime onset age and tissue cSL load controlled for tissue single-gene load (for hcSL network). Only age groups less than or equal to 40 years are considered for this analysis. Normal tissue GTEx gene expression data is used to compute the tissue cSL load.*

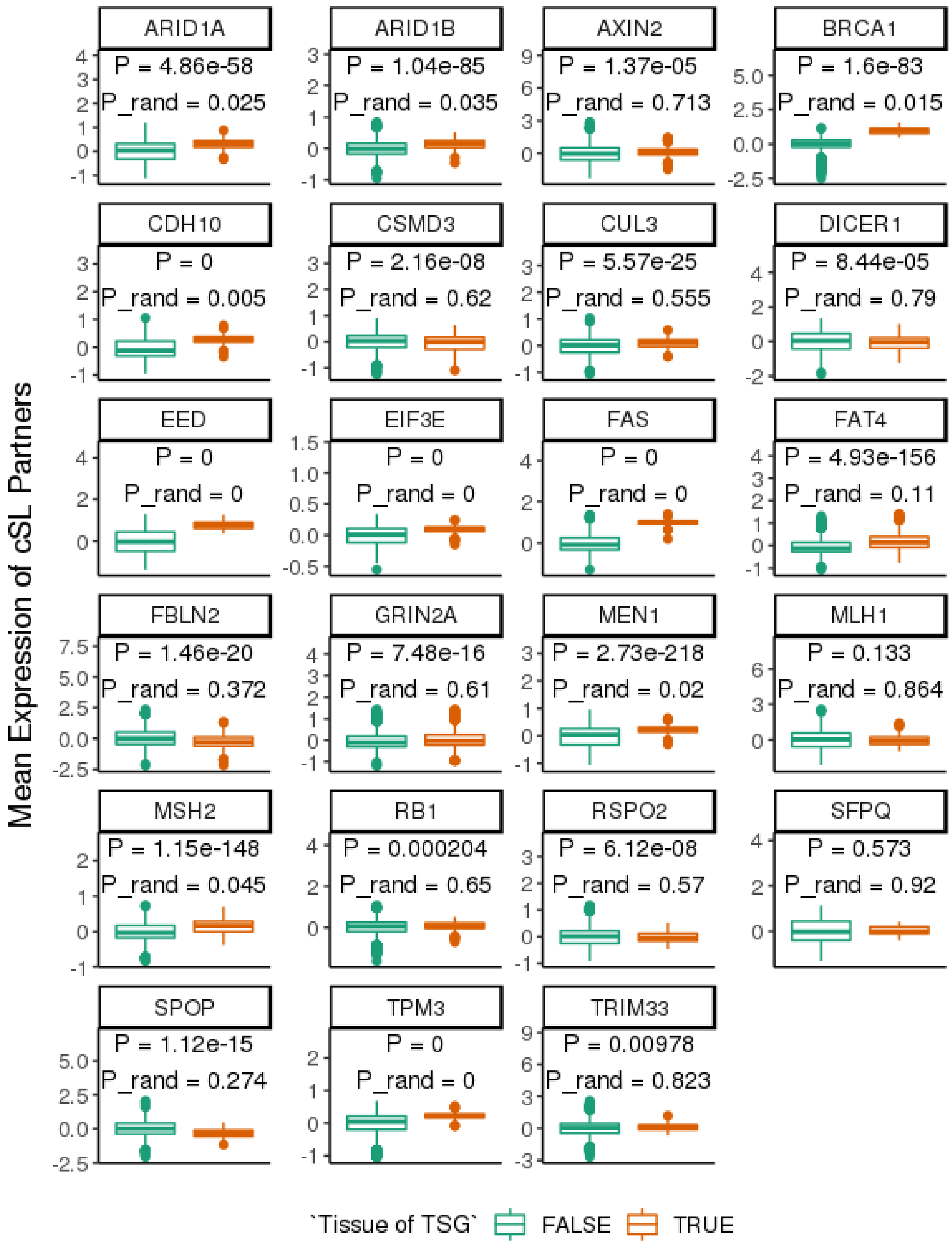


**Figure S8.** *The difference between the mean expression levels of the cSL partner genes of each tumor suppressor gene (TSG) in the tissue type(s) where the TSG is a known driver ("Tissue of TSG" is TRUE) and in the rest of the tissue types where the TSG is not an established driver ("Tissue of TSG" is FALSE). The expression levels are those from the GTEx normal tissue samples. Each panel is a tissue-specific TSG, and the tissue*

*types where they are established cancer drivers can be found in Table S6 (Note that no cSL partner genes were identified for most of the TSGs in Table S6, and we are analyzing only the TSGs with more than one cSL partners identified). The P values (top row) shown are from the linear models associated with the "Tissue of TSG" term as described in the main Methods, and the "P_rand" values (bottom row) are the empirical P values obtained from the random control tests described in the Supp. Notes. Where the P values are numerically extremely small, they are displayed as zero by the software.*

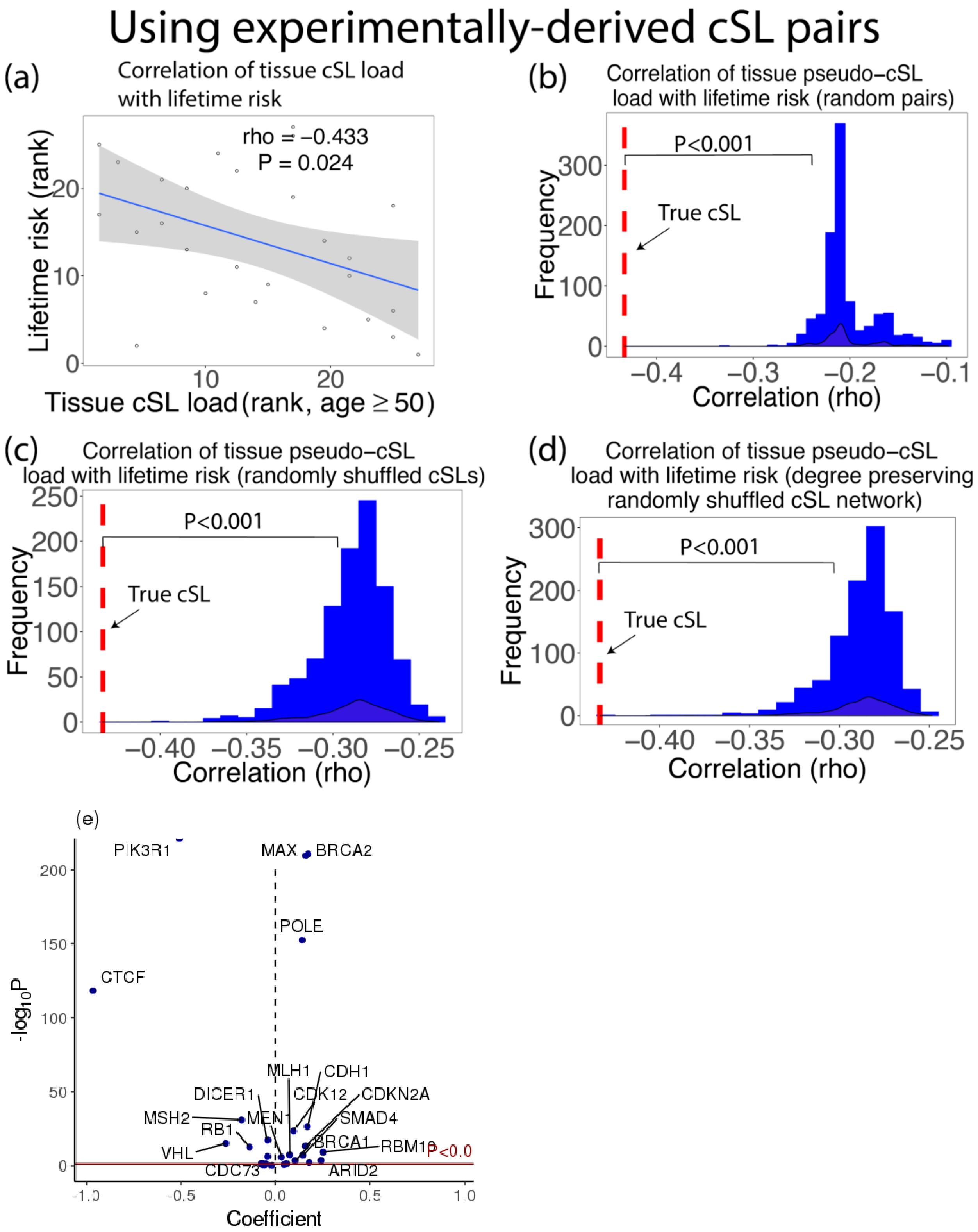


**Figure S9.** *(a) Spearman correlation (rho) between tissue cSL loads computed from experimentally identified cSL pairs with lifetime cancer risk. (b) Histogram showing*

*Spearman correlations of pseudo-cSL load (computed from random gene pairs) with cancer lifetime risk (no. of iterations = 1000). (c) Histogram showing Spearman correlations of pseudo-cSL load (computed from randomly shuffled experimentally identified cSL pairs) with cancer lifetime risk (no. of iterations = 1000). (d) Histogram showing Spearman correlations of pseudo-cSL load (computed from randomly shuffled experimentally identified cSL pairs) with cancer lifetime risk (no. of iterations = 1000). (d) Histogram showing Spearman correlations of pseudo-cSL load (computed from degree-preserving randomized experimentally identified cSL network) with cancer lifetime risk (no. of iterations = 1000). In subfigures (b,c,d) Correlation for cSL load (from experimentally identified cSLs -- true cSL) and cancer risk is shown in red. Randomization test shows that the correlation obtained from true cSL network is much more significant than those obtained from pseudo-cSL loads (P < 0.001). (e) For each tissue-specific tumor suppressor (TSG) gene Gi, the expression levels of its experimentally identified cSL partner genes in the tissue type(s) where gene Gi is a TSG were compared to those where gene Gi is not an established TSG, using GTEx normal tissue expression data. The volcano plot summarizes the result of comparison with linear models. Positive linear model coefficients (X-axis) mean that the expression levels of the cSL partner genes are on average higher in the tissue(s) where gene Gi is a TSG. Many cases have near-zero P values and are represented by points (half-dots) on the top border line of the plot. All TSGs with FDR < 0.05 are labeled.*

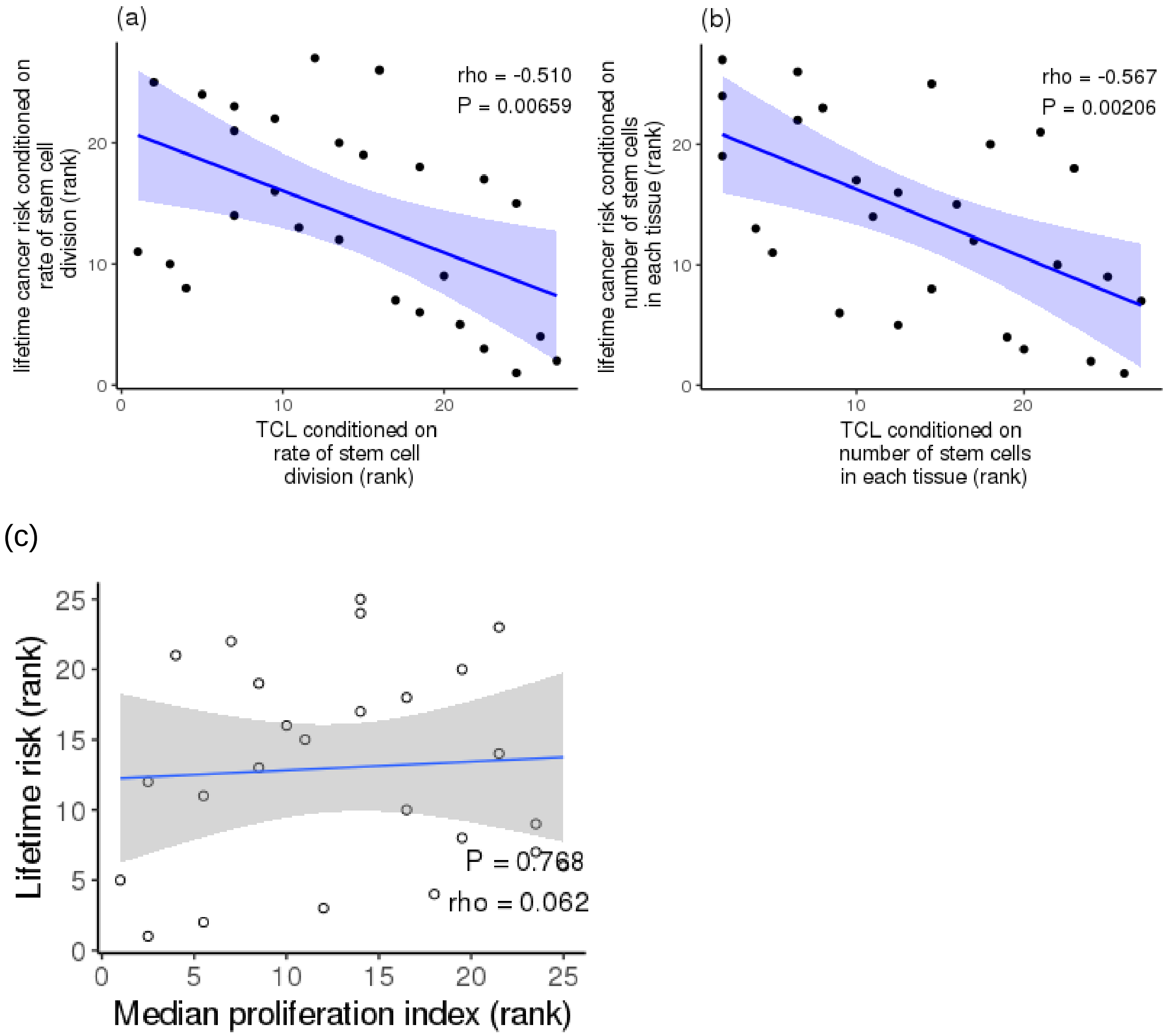


**Figure S10.** *Scatter plots showing partial Spearman correlations (rho) between lifetime cancer risk and tissue cSL load (TCL) conditioned on: (a) the rate of tissue stem cell division; (b) the number of stem cells residing in each normal human tissue. (c) A scatter plot showing the Spearman correlations (rho) between lifetime cancer risk and the median proliferation index for each type of the normal tissues from the GTEx dataset.*

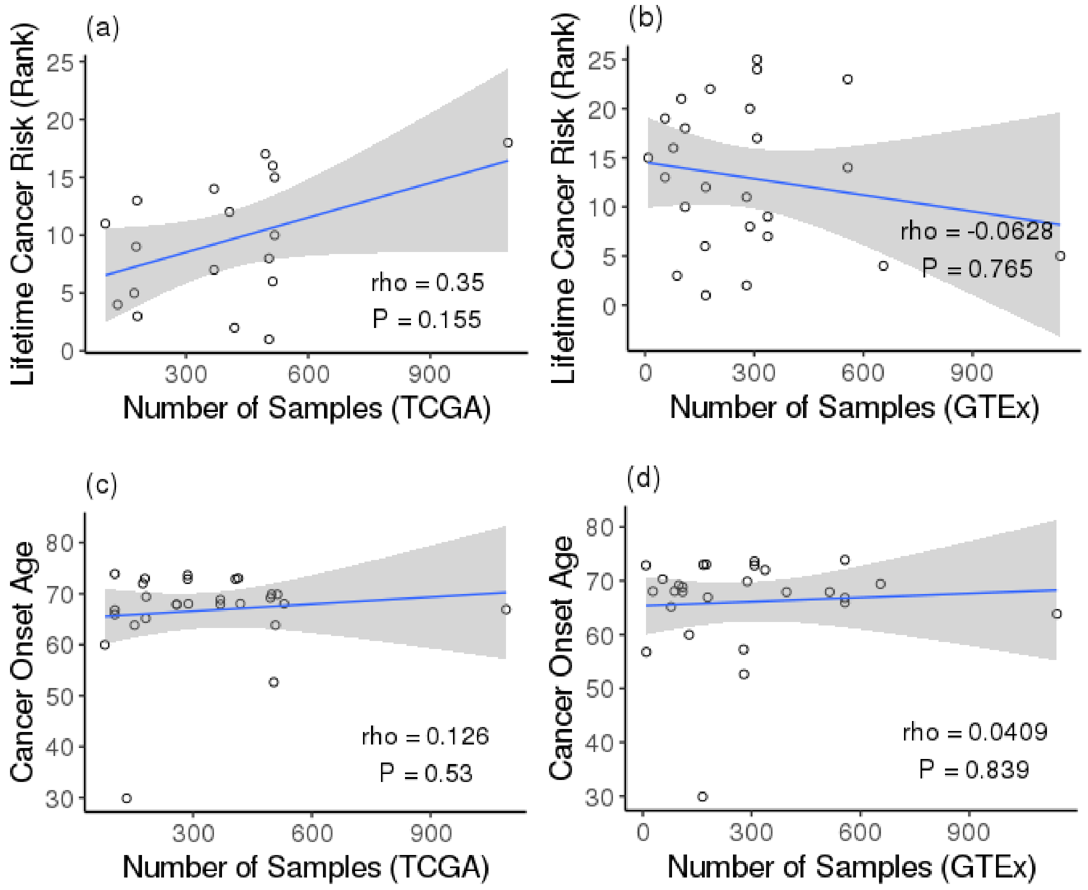


**Figure S11.** *The number of samples of each tissue in the TCGA or GTEx datasets is not a confounding factor for correlation with cancer risk or onset age. The four scatter plots show all the pairwise correlation between the number of TCGA or GTEx samples for each tissue type and lifetime cancer risk (ranked) or cancer onset age. Spearman's correlation coefficient and the corresponding P value are shown.*
